# Litter commensal bacteria can limit the horizontal gene transfer of antimicrobial resistance to *Salmonella* in chickens

**DOI:** 10.1101/2021.04.02.438293

**Authors:** Adelumola Oladeinde, Zaid Abdo, Benjamin Zwirzitz, Reed Woyda, Steven M. Lakin, Maximilian O. Press, Nelson A. Cox, Jesse C. Thomas, Torey Looft, Michael J. Rothrock, Gregory Zock, Jodie Plumblee Lawrence, Denice Cudnik, Casey Ritz, Samuel E. Aggrey, Ivan Liachko, Jonas R. Grove, Crystal Wiersma

**Affiliations:** U.S. National Poultry Research Center, USDA-ARS, Athens, GA, USA; Department of Microbiology, Immunology and Pathology, Colorado State University, Fort Collins, Colorado, USA; Institute of Food Science, University of Natural Resources and Life Sciences, Vienna, Austria; Phase Genomics Inc, Seattle, WA, 98109, USA; Office of National Programs, USDA-ARS, Beltsville, Maryland, USA; Division of STD Prevention, National Center for HIV/AIDS, Viral Hepatitis, STD and TB Prevention, Center for Disease and Control, Atlanta, Georgia, USA; National Animal Disease Center, USDA-ARS, Ames, IA, USA; Poultry Science Dept, University of Georgia, Athens, GA, USA

## Abstract

Fostering a ’balanced’ gut microbiome through the administration of beneficial microbes that can competitively exclude pathogens has gained a lot of attention and use in human and animal medicine. However, little is known about how microbes affect the horizontal gene transfer of antimicrobial resistance (AMR). To shed more light on this question, we challenged neonatal broiler chicks raised on reused broiler chicken litter – a complex environment made up of decomposing pine shavings, feces, uric acid, feathers, and feed, with *Salmonella* Heidelberg (*S*. Heidelberg), a model pathogen. Neonatal chicks challenged with *S*. Heidelberg and raised on reused litter were more resistant to *S*. Heidelberg cecal colonization than chicks grown on fresh litter. Furthermore, chicks grown on reused litter were at a lower risk of colonization with *S*. Heidelberg strains that encoded AMR on IncI1 plasmids. We used 16S rRNA gene sequencing and shotgun metagenomics to show that the major difference between chicks grown on fresh litter and reused litter was the microbiome harbored in the litter and ceca. The microbiome of reused litter samples was more uniform and enriched in functional pathways related to the biosynthesis of organic and antimicrobial molecules than fresh litter samples. We found that *E. coli* was the main reservoir of plasmids encoding AMR and that the IncI1 plasmid was maintained at a significantly lower copy per cell in reused litter compared to fresh litter. These findings support the notion that commensal bacteria play an integral role in the horizontal transfer of plasmids encoding AMR to pathogens like *Salmonella*.

**Importance/Significance:** Antimicrobial resistance spread is a worldwide health challenge, stemming in large part, from the ability of microorganisms to share their genetic material through horizontal gene transfer. To address this issue, many countries and international organization have adopted a One health approach to curtail the proliferation of antimicrobial resistant bacteria. This includes the removal and reduction of antibiotics used in food animal production and the development of alternatives to antibiotics. However, there is still a significant knowledge gap in our understanding of how resistance spreads in the absence of antibiotic selection and the role commensal bacteria play in reducing antibiotic resistance transfer. In this study, we show that commensal bacteria play a key role in reducing the horizontal gene transfer of antibiotic resistance to *Salmonella* and provide the identity of the bacterial species that potentially perform this function in broiler chickens and also postulate the mechanism involved.

## Introduction

Horizontal gene transfer (HGT) is recognized as the main mechanism by which bacteria acquire antimicrobial resistance (AMR) and exposure to antibiotics has been shown to drive antimicrobial resistance gene (ARG) transfer [1]. Consequently, the rise in AMR in bacteria from hospital settings has been linked to the overuse of antibiotics in humans and their use in food animal production [2, 3]. These public health concerns have led to a reduction in antibiotics used for raising food animals [4, 5], including a ban on antibiotics’ use in Europe [6]. We previously showed that neonatal broiler chicks challenged with a nalidixic acid resistant *S*. Heidelberg strain and raised antibiotic-free on fresh litter composed of pine shavings were colonized at a high rate with *S*. Heidelberg strains that harbored IncI1 plasmids encoding AMR [7]. We selected *S*. Heidelberg as the model pathogen because of its promiscuity to plasmids carrying AMR [8, 9].

There is limited research on if and how commensal bacteria reduce AMR in food-borne pathogens [10–16], such as *Salmonella enterica*, and the role it plays in AMR reduction and transfer. Therefore, the goal of this study was to determine the role commensal bacteria play in limiting *Salmonella enterica* from acquiring AMR and to provide information on the bacterial species and mechanism involved. To do this, we compared the dynamics of AMR transfer in neonatal broiler chicks raised on fresh litter to chicks raised on reused litter. We chose reused litter because it is a complex environment made up of decomposing plant-based bedding (e.g., wood shavings, sawdust and rice or peanut hulls) mixed with chicken feces, uric acid, feathers, feed, insects, and other broiler-sourced materials. Therefore, reused litter carries a unique and complex population of bacteria, fungi, and viruses [17–19], interacting with various forms of eukaryotes, making it a suitable environment to study competitive exclusion. Broiler chickens are commonly raised on litter but how litter is managed differs between countries, producers, and farmers. For instance, the practice of reusing litter over multiple flocks of broiler chickens is a widespread practice in the US and Brazil while Canada [20] and Europe [21] recommend fresh litter bedding for every flock. One argument against litter reuse is that it harbors pathogenic bacteria that can be transferred to the next flock [22]. Contrastingly, proponents of litter reuse argue that it confers competitive exclusion against pathogens when effectively managed and it is cost-effective [23–26]. Beyond broiler chicken production, competitive exclusion by commensal bacteria has received enormous attention in the 20th century, resulting in the bloom of commercially marketed probiotics [27] and the underlying theory behind the application of fecal microbiota transplantation in human medicine [28].

## Results

### *S*. Heidelberg abundance and prevalence in ceca and litter

We confirmed that neonatal chicks were *Salmonella*-free by testing the chick pads used for transportation from the hatchery for *Salmonella*. None of the chick pads were positive for *Salmonella.* The reused litter for this study was confirmed to be *Salmonella*- free in earlier studies [29, 30]. To determine if the microbiome offers protection against *Salmonella* colonization, we challenged neonatal broiler chicks with a nalidixic resistant (nal^R^) strain of *S*. Heidelberg and raised the chicks on either fresh pine shavings (here after referred to as fresh litter) (n = 75) or reused litter (litter previously used to raise three flocks of broiler chickens) (n =75). Chicks were challenged either through oral gavage (n = 25), cloacal inoculation (n =25), or by the seeder method (i.e., a few chicks [n = 5] were challenged orally and comingled with unchallenged chicks [n = 20]) [7]. Unchallenged chicks on fresh (n =25) and reused litter (n =25) were used as controls. Broiler chicks on fresh and reused litter were housed separately for 14 days (four separate pens for each treatment in a house) and chicks were not administered any medication or antibiotics for the duration of the study.

Afterwards, we determined the concentration of nal^R^ *S*. Heidelberg in the ceca and litter of chicks 14 days after they were challenged. Average nal^R^ *S*. Heidelberg concentration in the ceca (n=10 for oral and cloacal, n=15 for seeder) was not significantly different between chicks raised on fresh litter compared to reused litter (W-statistic, *P*. value; oral = 41, 0.15; cloacal= 41, 0.96; seeder = 5, 1.00) (Fig.1a). However, the ceca of chicks raised on fresh litter were more likely to be positive for nal^R^ *S*. Heidelberg (X_2_= 16.07, df = 2, *P* = 0.0003), than chicks on reused litter. The percentage of cecal samples positive for nal^R^ *S*. Heidelberg was 100%, 100% and 50%, respectively for oral, cloacal and seeder chicks raised on fresh litter, compared to 80%, 100% and 33.3 %, for chicks raised on reused litter. For the seeder treatment on fresh litter, one seeder was lost due to premature mortality and three of the four seeders were positive for nal^R^ *S*. Heidelberg, while the ceca of four of the ten uninoculated contact chicks were positive. In contrast, the ceca of the five orally challenged seeders raised on reused litter were positive for nal^R^ *S*. Heidelberg, but all contact chicks were negative (n = 10).

**Fig 1.**
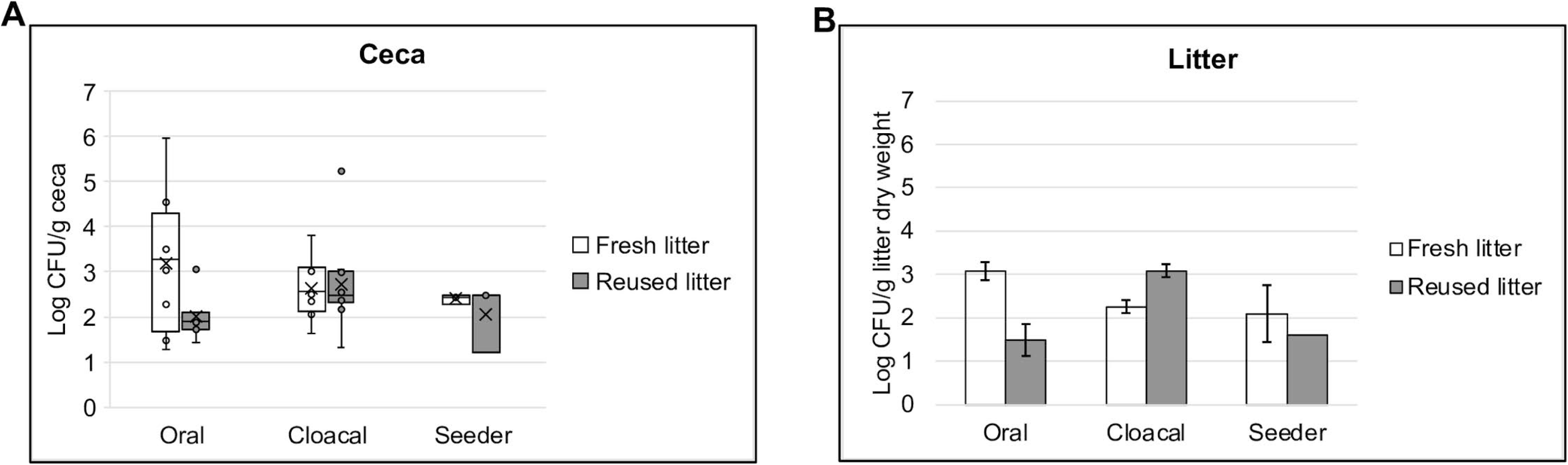
***S*. Heidelberg concentration in the ceca and litter of broiler chicks**. Box plot of nal^R^ *S*. Heidelberg concentration in the (**a**) ceca (n=10 for oral and cloacal, n=15 for seeder) (**b**) and litter (n = 2 subsamples from pooled litter) of chicks raised on fresh or reused litter (*P >* 0.05; Wilcoxon signed-rank test). Cecal and litter samples were collected 14 days after challenging chicks with nal^R^ *S*. Heidelberg through oral gavage, cloacal inoculation or through the seeder method.

For the litter, oral and seeder treatments carried higher levels of nal^R^ *S*. Heidelberg in fresh litter compared to reused litter, while cloacally inoculated chicks had higher levels in reused litter compared to fresh litter (Fig.1b). All litter samples from challenged chicks were positive for nal^R^ *S*. Heidelberg by direct culture or after enriching in buffered peptone water (BPW), except for one reused litter sample from the seeder treatment that was negative. For unchallenged chicks used as controls, one litter sample from fresh and reused litter tested positive after enrichment in BPW. These data suggests that litter age/type did not have a significant effect on the abundance of nal^R^ *S*. Heidelberg in the ceca, but chicks raised on reused litter (66%) had a lower *Salmonella* positivity rate compared to chicks on fresh litter (79%).

### Broiler litter age/type affected the horizontal transfer of AMR

Next, we questioned if the litter plays a role in the HGT of AMR. First, we performed antibiotic susceptibility testing (AST) on *S*. Heidelberg isolates recovered from the ceca and litter of broiler chicks raised on fresh litter (n=158) and reused litter (n=141). As expected, all *S*. Heidelberg isolates were resistant to nalidixic acid (Fig. 2). Thirty-one percent of isolates from the ceca (37/118) and 37.5 % from the litter (15/40) of chicks on fresh litter acquired resistance to gentamicin, tetracycline, and streptomycin [7] (Fig. 2a). Contrastingly, 3% of *S*. Heidelberg isolates from the ceca of chicks on reused litter acquired resistance to either cefoxitin (1/114) or tetracycline (2/114), while ∼18% of isolates from reused litter acquired resistance to either ciprofloxacin (1/27), tetracycline (2/27) or streptomycin (2/27) (Fig. 2a). The percentage of *S*. Heidelberg isolates that acquired AMR in fresh litter differed by the route of inoculation [7]. For instance, 40% of isolates from orally challenged chicks acquired AMR compared to 24% for cloacal [7].

**Fig 2.**
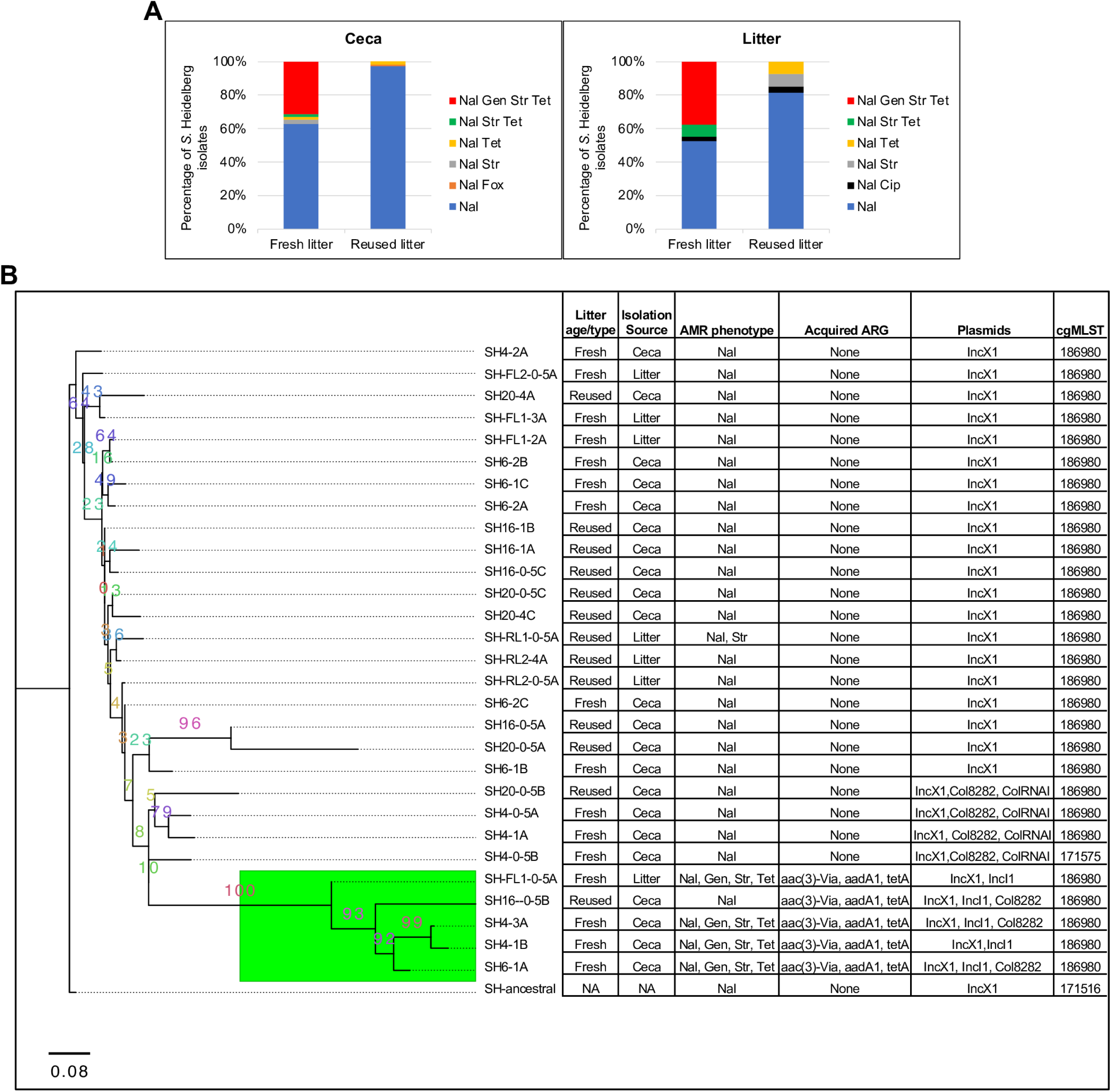
**Broiler chicks raised on reused litter carried lower levels of antimicrobial resistant *S*. Heidelberg.** (**a**) Distribution of antibiotic resistance acquired by nal^R^ S. Heidelberg isolates from the ceca and litter of chicks on fresh (n=158) and reused litter (n=141). (**b**) Maximum likelihood tree constructed using accessory genes present in *S*. Heidelberg isolates (*n* = 30) recovered from the ceca and litter of cloacally inoculated chicks raised on fresh and reused litter. The GTR model of nucleotide substitution and the GAMMA model of rate heterogeneity were used for sequence evolution prediction. Numbers shown next to the branches represent the percentage of replicate trees where associated isolates cluster together based on ∼100 bootstrap replicates. All *S*. Heidelberg strains were assembled using Illumina short reads except SH4-3A and SH- ancestral that were assembled by combining Illumina short reads with PacBio or MinION long reads. The tree was rooted with the ancestral *S*. Heidelberg strain (SH-ancestral, Genbank accession no: CP066851). Green colored subclade shows strains that harbored IncI1-pST26. All isolates are expected to be resistant to Nalidixic acid. (Nal, nalidixic acid; Gen, gentamicin; Str, streptomycin; Tet, tetracycline; Fox, Cefoxitin; Cip, Ciprofloxacin)

We determined if *S*. Heidelberg isolates recovered from chicks raised on fresh or reused litter differed in their core genome or if they acquired new mobile elements like plasmids. To answer this question, we performed whole genome sequence (WGS) analysis on *S*. Heidelberg isolates (n=29) recovered from the ceca and litter of chicks that were cloacally challenged. We chose cloacal samples because only this route of challenge resulted in 100% colonization of the ceca of chicks raised on fresh and reused litter. There was no difference in the core genome multilocus sequence types (cgMLST) of evolved *S*. Heidelberg isolates from fresh and reused litter, however, 28 of the 29 evolved isolates had a different cgMLST from the ancestral nal^R^ *S*. Heidelberg (Fig. 2b). We used the genomes sequenced to construct a maximum likelihood (ML) tree based on the pangenome and mutations of *S*. Heidelberg strains recovered. The core genome (genes present in ≥ 95% of the strains) and accessory genome (genes present in < 95% of the strains) was composed of 4,373 and 356 genes, respectively. After removing 10 isolates with identical mutations, a total of 118 informative sites (SNPs and indels) was used for ML SNP tree construction.

The three constructed ML trees did not group isolates by litter age/type (fresh versus reused) (Fig. 2b, Fig. S1a and b), but *S*. Heidelberg isolates that acquired IncI1 plasmids (plasmid MLST 26; hereafter referred to as IncI1-pST26) (n=5) formed a separate clade on the accessory genome tree (Fig. 2b). The IncI1-pST26 plasmid harbored the aminoglycoside (*aadA1*), tetracycline (*tetA*) and mercury (*mer* operon) resistance genes [7] (Fig. S2). Isolates from the ceca (n=3) and litter (n=1) of chicks on fresh litter that harbored IncI1-pST26 displayed the antibiotic resistance phenotype predicted by WGS (Fig. 2b). Contrastingly, one isolate from reused litter carried IncI1-pST26 but was found to be susceptible to all antibiotics tested except for the ancestral nalidixic resistance. These results indicated that chicks grown on reused litter are less likely to carry *S*. Heidelberg isolates harboring AMR compared to chicks on fresh litter.

### The microbiome of neonatal chicks differed for fresh and reused litter

We used 16S rRNA gene sequencing to examine the differences in the microbiome between neonatal chicks raised on fresh and reused litter. The beta diversity of the cecal (n = 59) and litter (n = 22) microbiome of chicks was significantly different between fresh and reused litter (ceca, *P* = 0.0002; litter, *P* = 0.0002) (Fig. 3a). Furthermore, the route of inoculation used for *S*. Heidelberg affected the beta diversity of the ceca and litter for fresh and reused litter (*P* < 0.05). For example, chicks challenged orally (n = 10) or through the cloaca (n =10) and raised on fresh litter harbored a significantly different beta diversity than uninoculated chicks (n = 10) in the ceca (*P* < 0.002) (Fig. 3a). Likewise, the beta diversity of the ceca of chicks challenged orally (n = 10) differed from cloacal (n = 9) and uninoculated controls (n = 10) for reused litter (*P* < 0.05). In the litter, pair wise comparisons between routes of inoculation were not significant (*P* > 0.05) for beta diversity. Nevertheless, fresh litter samples (n = 3) from orally challenged chicks clustered next to litter samples (n = 3) from seeder treatments, while litter samples (n = 2) from cloacally inoculated chicks were different from oral and seeder treatments (Fig. 3a). For reused litter samples, litter from orally (n = 3) and cloacally (n = 3) challenged chicks clustered together, while litter (n = 2) from seeder treatments was similar to litter samples (n = 3) from control chicks. Lastly, the bacterial community structure (assessed using beta dispersion) of reused litter was less variable than fresh litter (*P* = 0.001), however there was no difference in variability for the ceca (*P* = 0.15). Moreover, the route of challenge affected the bacterial community beta dispersion in the ceca and litter of chicks on fresh and reused litter (*P* = 0.001).

**Fig 3.**
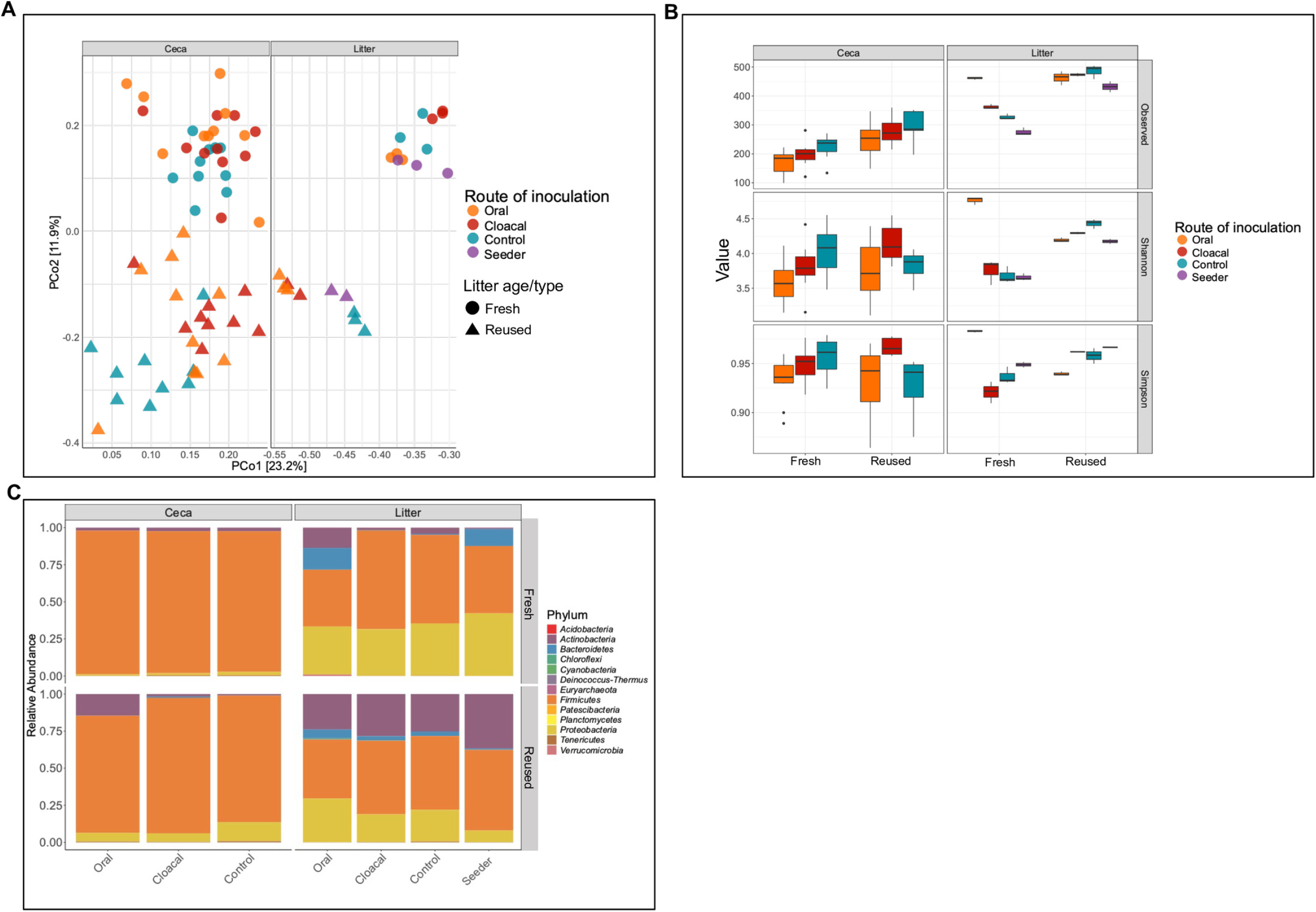
**Reused litter samples harbored a uniform and diverse microbiome than fresh litter samples.** (**a**) Principal coordinates analysis of Bray-Curtis distances based on 16S rRNA gene libraries obtained from ceca and litter samples. Each point is values from individual libraries with colors denoting the route used for *S*. Heidelberg challenge for respective samples. (b) Average alpha diversity indices of rarefied ceca and litter samples grouped by the litter age/type and the route of *S*. Heidelberg challenge. Boxes indicate the interquartile range (75th to 25th) of the data. Whiskers extend to the most extreme value within 1.5 * interquartile range and dots represent outliers beyond that range. Black bar represents the median. (**c)** Phylum-level classification of 16S rRNA gene sequence reads in each ceca and litter sample grouped by the route of *S*. Heidelberg challenge. 16S rRNA gene sequencing was performed on the ceca (59 chicks from fresh litter (n = 30) and reused litter (n = 29)) and litter (n = 22 (11 samples each from fresh and reused litter)).

The alpha diversity (i.e., the number of amplicon sequence variants (ASVs)) of the microbiome present in the ceca and litter of chicks on reused litter was higher for the Observed species measure of diversity (ceca, *P* < 0.001; litter, *P* = 0.001) but not for Shannon (ceca, *P* = 0.25; litter, *P* = 0.05) and Simpson (ceca, *P* = 0.97; litter, *P* = 0.28) indices. The route of *S*. Heidelberg challenge affected the alpha diversity of the cecal and litter microbiome (*P* < 0.05) (Fig. 3b). The alpha diversity of the ceca of challenged and control chicks raised on reused litter was higher than chicks grown on fresh litter for Observed species measure of diversity (*P* < 0.01). Cloacal challenged chicks raised on reused litter had higher alpha diversity in their ceca compared to chicks on fresh litter for Shannon and Simpson indices (*P* < 0.05). Uninoculated control chicks on fresh litter had higher alpha diversity in their ceca than control chicks on reused litter for Simpson index, (Fig. 3b). The litter of cloacal, seeder, and control chicks on reused litter had higher alpha diversity than the litter of chicks on fresh litter for Shannon measure of diversity (*P* < 0.05) (Fig. 3b). In contrast, the litter of orally challenged chicks on fresh litter had higher alpha diversity than the litter of gavaged chicks on reused litter for Shannon index (*P* = 0.0001). There was no significant difference in alpha diversity between fresh and reused litter for Observed and Simpson indices (*P* > 0.05).

On the taxonomic level, orally challenged chicks on reused litter carried higher abundance of Actinobacteria in the ceca compared to chicks on fresh litter (*P* = 0.003), while uninoculated control chicks on fresh litter had higher relative abundance of Actinobacteria in the ceca compared to control chicks on reused litter (*P* = 0.004) (Fig. 3c). Proteobacteria was higher in the ceca of oral, cloacal and control chicks on reused litter compared to chicks on fresh litter (*P* < 0.01). Firmicutes had higher relative abundance in the ceca of chicks on fresh litter compared to reused litter for all treatments (*P* < 0.01) while Bacteroidetes had higher relative abundance in the ceca of cloacal and control chicks on reused litter compared to chicks on fresh litter (*P* < 0.05) (Fig. 3c). For the litter, Actinobacteria was higher in reused litter compared to fresh litter (*P* < 0.05) while Proteobacteria and Bacteroidetes were higher in fresh litter compared to reused litter (*P* < 0.05). There was no significant difference in phyla abundance between routes of inoculation (*P* > 0.05) for the litter. These results shows that the microbiome of chicks raised on fresh litter differed from chicks grown on reused litter, and reused litter cecal and litter samples of challenged chicks were more uniform than fresh litter samples.

### *S.* Heidelberg challenge modulated the gut microbiome

We determined that the core microbiome (i.e., ASVs > 0.1% relative abundance and present in at least 80 % of samples) of fresh litter differs from reused litter (Table S1) and found that *S*. Heidelberg challenge perturbed the cecal and litter microbiome of broiler chicks (Fig. 4). For example, Enterobacteriaceae (*Klebsiella pneumonaie* and *Proteus mirabilis*) were part of the core microbiome in fresh litter, while they were not part of the core microbiome of reused litter (Table S1). Contrastingly, Actinobacteria (23 ASVs) was part of the core microbiome of reused litter, while only one ASV classified as Actinobacteria was determined to be a core member of fresh litter (Table S1). Unsurprisingly, the bacterial ASVs that increased or decreased in abundance after *S*. Heidelberg challenge differed between chicks on fresh and reused litter (Fig. 4a and b).

**Fig 4.**
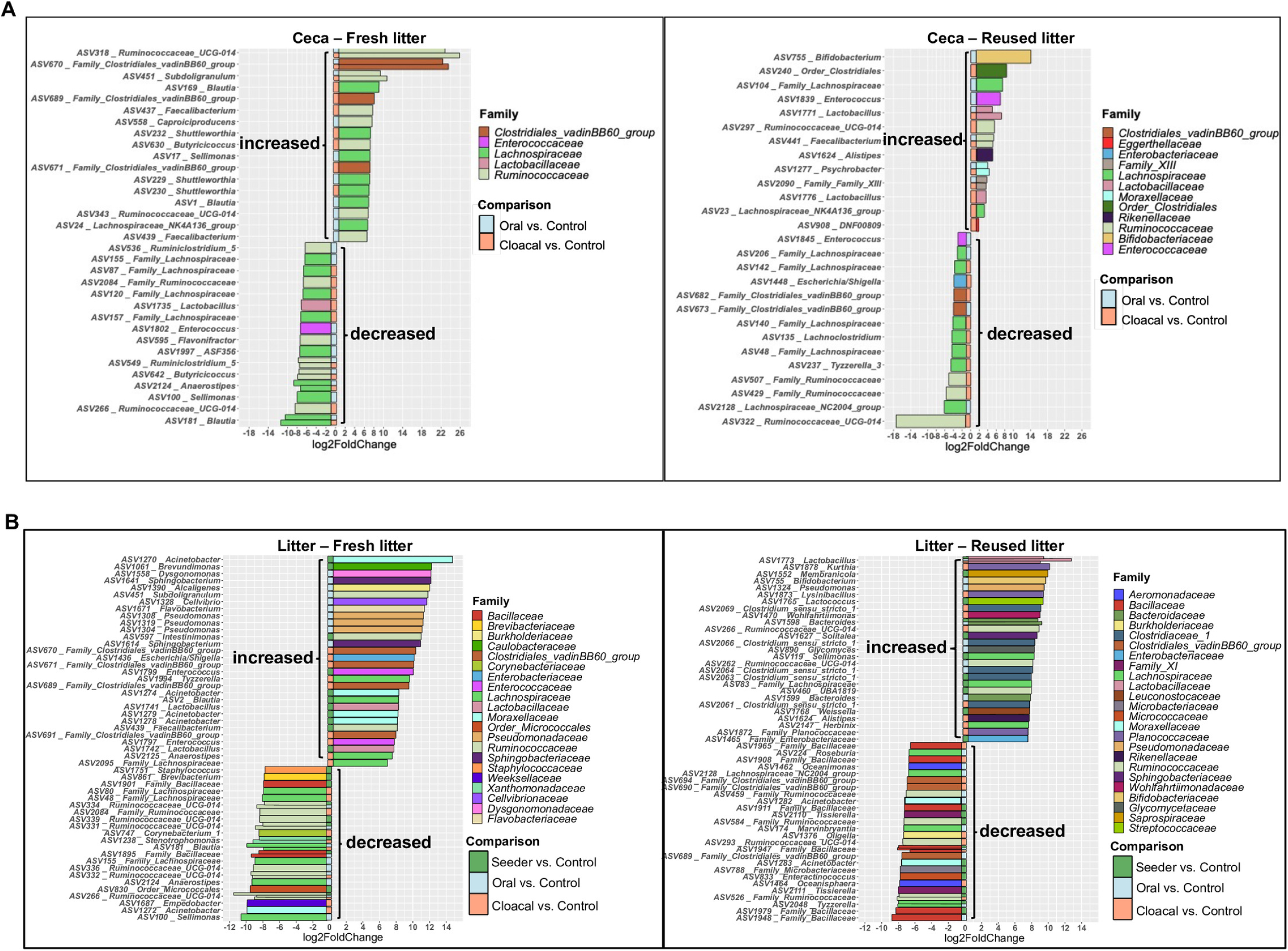
***S*. Heidelberg challenge modulated the microbiome of the ceca and litter of broiler chicks.** Plots of amplicon sequence variants (ASVs) that were significantly differentially abundant (*p*_adj_ < .05) in the (**a**) ceca and (**b**) litter of challenged chicks on fresh and reused litter compared to uninoculated controls. Significant ASVs are plotted individually and colored according to their family-level classification. The route of *S*. Heidelberg challenge is shown on the edge of each plot as colored squares.

To determine the ASVs that were modulated by *S*. Heidelberg challenge (i.e., ASVs that increased or decreased after challenge), we compared the relative abundance of ASVs in the ceca and litter of challenged chicks to uninoculated control chicks (Fig. 4). In the ceca of chicks on fresh litter, the ASVs significantly modulated were assigned to five bacterial families (Fig. 4a). Three ASVs matching family Clostridiales vadinBB6 group, nine ASVs classified as Ruminococcaceae and seven ASVs classified as Lachnospiraceae increased in abundance in challenged chicks compared to control chicks. Contrastingly, ASVs classified as Lactobacillaceae, Enterococcaceae, Ruminococcaceae (n=8) and Lachnospiraceae (n=10) decreased in abundance in challenged chicks compared to control chicks (Fig. 4a).

For the ceca of chicks on reused litter, the ASVs modulated were assigned to twelve bacterial families. Amplicon sequence variants matching Bifidobacteriaceae, order Clostridiales, Moraxellaceae, Rikenellaceae, Eggerthellaceae, Enterococcaceae, Family XIII, Lactobacillaceae (n=2), Ruminococcaceae (n=2) and Lachnospiraceae (n=2) increased in abundance in challenged chicks compared to control chicks. In contrast, ASVs classified as Enterobacteriaceae, Enterococcaceae, Clostridiales vadinBB6 group (n=2), Ruminococcaceae (n=3) and Lachnospiraceae (n=7) decreased in abundance in challenged chicks compared to control chicks (Fig. 4a).

For the litter, the ASVs that were significantly modulated in fresh litter were assigned to twenty-one bacterial families (Fig. 4b). The ASVs that increased in abundance in fresh litter of challenged chicks belonged to Caulobacteraceae, Dysgonomonadaceae, Burkholderiaceae, Flavobacteriaceae, Cellvibrionaceae, Enterobacteriaceae, Enterococcaceae (n=2), Lactobacillaceae (n=2), Sphingobacteriaceae (n=2), Pseudomonadaceae (n=3), Ruminococcaceae (n=3), Moraxellaceae (n=4), Clostridiales vadinBB6 group (n=4), and Lachnospiraceae (n=4). Conversely, ASVs assigned to Staphylococcaceae, Brevibacteriaceae, order Micrococcales, Corynebacteriaceae, Xanthomonadaceae, Weeksellaceae, Bacillaceae (n=2), Ruminococcaceae (n=6) and Lachnospiraceae (n=6) decreased in abundance in the litter of challenged chicks compared to control chicks (Fig 4b).

Amplicon sequence variants modulated in reused litter after challenge were grouped into twenty-four bacterial families. The ASVs that increased in abundance in reused litter of challenged chicks were assigned to Lactobacillaceae, Saprospiraceae, Bifidobacteriaceae, Rikenellaceae, Pseudomonadaceae, Sphingobacteriaceae, Leuconostocaceae, Glycomycetaceae, Streptococcaceae, Enterobacteriaceae, Wohlfahrtiimonadaceae, Bacteroides (n=2), Ruminococcaceae (n=3), Planococcaceae (n=3), Lachnospiraceae (n=3) and Clostridiaceae 1 (n=5). The ASVs that decreased in the reused litter of challenged chicks were classified as Burkholderiaceae, Microbacteriaceae, Micrococcaceae, Moraxellaceae (n=2), Wohlfahrtiimonadaceae (n=2), Aeromonadaceae (n=3), Clostridiales vadinBB6 group (n=3), Ruminococcaceae (n=4), Lachnospiraceae (n=4) and Bacillaceae (n=6),

Not all bacterial families were modulated by the cloacal route of *S*. Heidelberg challenge. For instance, Enterococcaceae was only modulated by oral challenge in the ceca of chicks grown on fresh and reused litter (Fig. 4a). Similarly, Pseudomonadaceae was only modulated in the litter of chicks that were orally challenged (Fig. 4b). Together, these results show that the litter microbiome modulated by *S*. Heidelberg challenge differed between chicks on fresh litter and reused litter, and the route of challenge affected the ASVs that increased or decreased in abundance.

### Enterobacteriaceae and Clostridiales were the major bacterial hosts for AMR

We used Hi-C metagenomics to explore the bacterial reservoir of AMR in the cecal microbiome of 2 cecal samples each from cloacally challenged chicks on fresh and reused litter. We previously used this approach to identify *E. coli* as the main bacterial reservoir of transferable plasmids in fresh litter [7]. Here, we extended our analysis to cecal samples from reused litter. The average assembly size and number of contigs for the four samples was 1,238,395,273±167,820 bp and 1,797,423±281,253, respectively. The number of metagenome assembled genomes (MAGs) found in cecal samples from fresh and reused litter was 456±422. We searched the MAGs for sequences matching plasmid incompatibility (*inc*) groups and replicons available on the PlasmidFinder database. The *inc* groups found matched common Enterobacteriaceae plasmids including IncF, IncI, IncX, IncH, IncY, IncB/O/K/Z, p0111 and Col-like plasmids (Table 1). Three plasmid replicons were found for Gram-positive bacteria including *rep2*, *rep18b* and *repA* (Table 1). The low representation of plasmids is expected, since the PlasmidFinder database is much more comprehensive for Enterobacteriaceae than other bacterial species, and within the well-studied Proteobacteria most plasmids cannot be typed [31].

**Table 1.**
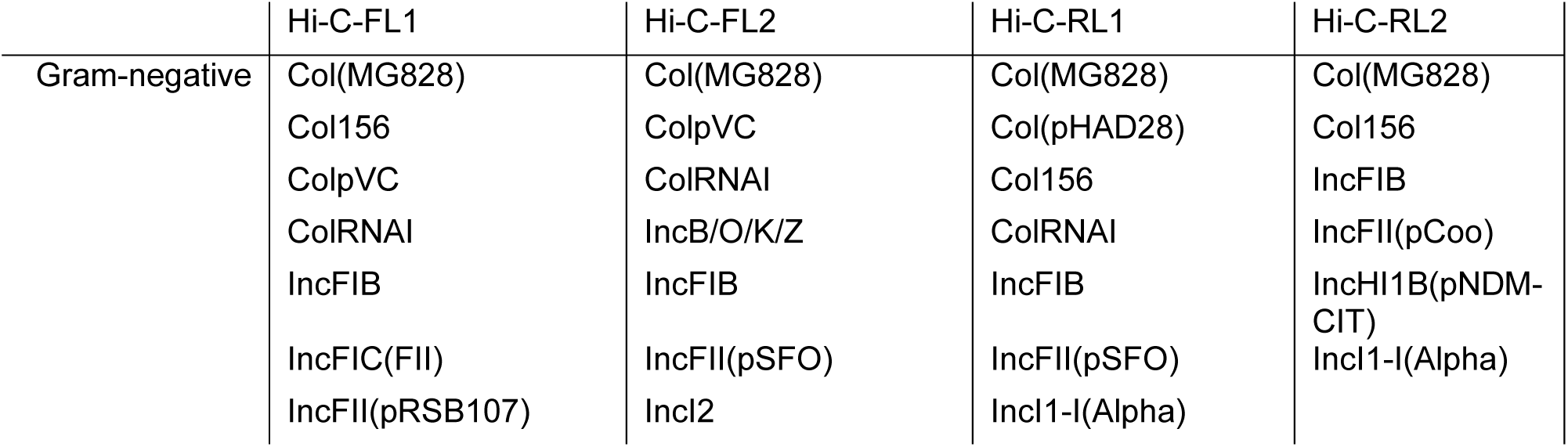
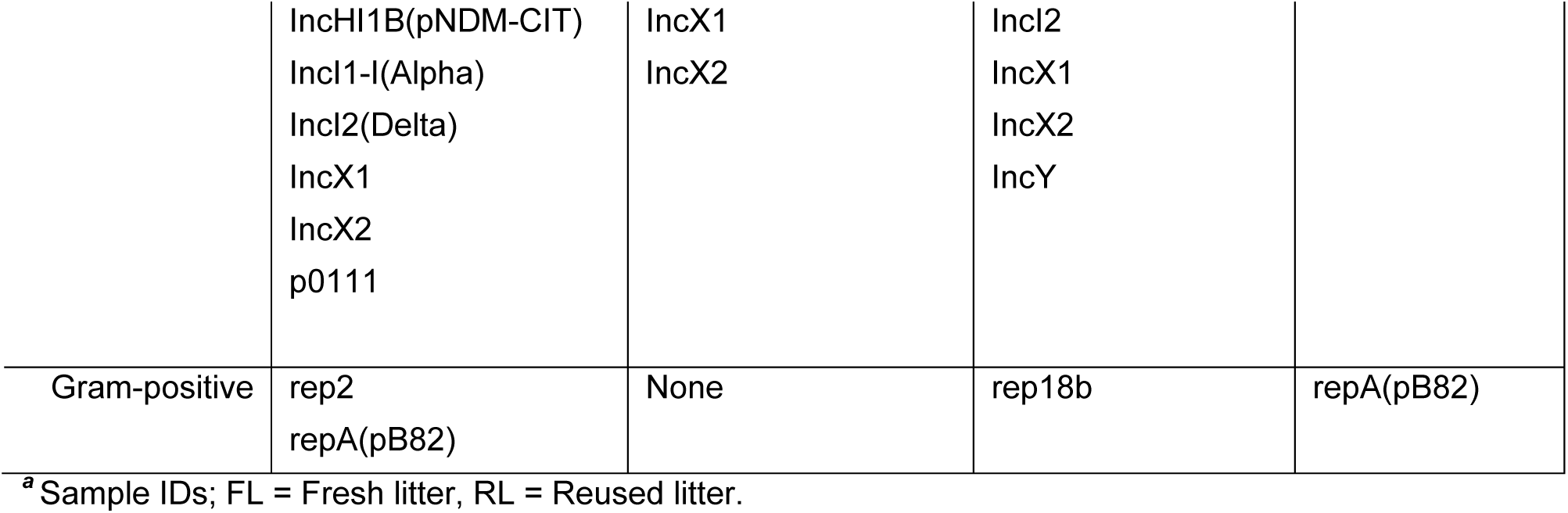
Plasmid incompatibility groups and replicons found in cecal metagenome assembled genomes using PlasmidFinder**^a^**.

Next, we used proximity-ligation to find the bacterial host of ARGs and to determine if they are encoded on plasmids or the chromosome. One hundred ARGs were found in the MAGs and the majority (∼ 50%) were harbored by family Enterobacteriaceae (Fig. 5). *Escherichia coli* was the only Enterobacteriaceae found to be the putative host for these ARGs in fresh and reused litter. Members of the order Clostridiales were the putative hosts for 27% of ARGs, while unclassified bacterial species (n = 6) and members of the phyla Firmicutes (n = 9), Bacteriodetes (n = 3) and Actinobacteria (n = 1) were the hosts for the remaining ARGs. Fifty-four percent of the ARGs were found to be encoded on the chromosome while 46% were found on plasmids. One cecal sample from fresh litter (Fig. 5) harbored *E. coli* MAGs that encoded silver and copper resistance genes (*silABCFRS* and *pcoABCDRS*) on the chromosome, while virulence genes (*iroBCDEN*) were found on plasmids. A cecal sample from a chick raised on reused litter harbored *E. coli* MAGs that encoded ampicillin (*bla_TEM1B_*), tetracycline (*tetA*), and mercury resistance (*merCPRT*) genes on plasmids (Fig. 5). This result suggests that the microbiome of chicks on fresh and reused litter harbored AMR, and Enterobacteriaceae and Clostridiales were the putative bacterial hosts of the ARGs.

**Fig 5.**
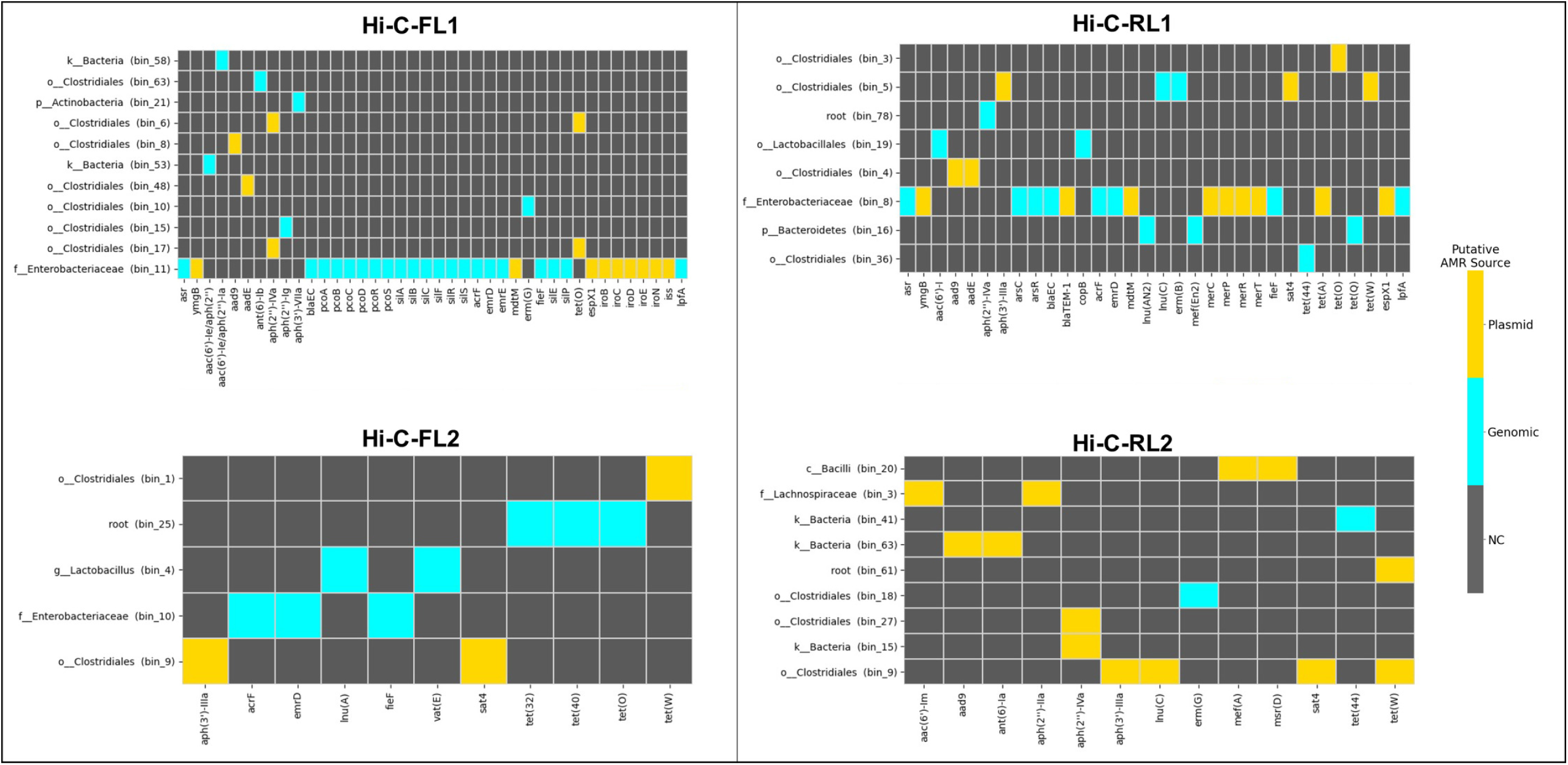
**Enterobacteriaceae and Clostridiales were the bacterial hosts of AMR in broiler chicks**. AMR gene hosts found by Hi-C contacts. AMR genes (horizontal axis of heatmap) associated with metagenome-assembled genomes (MAGs) present in cecal samples from chicks on fresh litter (Hi-C-FL1 and Hi-C-FL2) and reused litter (Hi-C-RL1 and Hi-C-RL2). MAGs and AMR gene sources were derived from Hi-C based deconvolution of the metagenome assembly and placed into a bacterial phylogeny using Mash and CheckM by the ProxiMeta platform. The MAG assigned to family Enterobacteriaceae in each sample matches most closely to an *Escherichia coli* genome. (Legend: Gold - AMR genes that are associated with plasmid sequences on the same contigs, Cyan – AMR genes that are likely to be genomically integrated, Black – no contact above statistical thresholds)

### *E. coli* isolates harbored AMR on plasmids and genomic islands

In our earlier work, we found that *E. coli* MAGs were the primary hosts of IncI1 by retrospectively screening the cecal contents of chicks on fresh litter for *E. coli* isolates [7]. Using this approach, we confirmed that the IncI1-pST26 plasmid acquired by *S*. Heidelberg was identical to the IncI1-pST26 plasmid present in *E. coli* strains (Fig. S2). Here, we performed AST and WGS on additional *E. coli* isolates recovered from the ceca of chicks used for Hi-C metagenomics (Table S2) and a pangenome analysis was done on nineteen of them. The isolates were randomly selected from either CHROMagar™ plates supplemented with or without gentamicin and tetracycline.

The inferred phylogeny of the *E. coli* strains using their core (n = 3385) and accessory genes (n = 6024) partially grouped the isolates by the litter used for raising the chicken host (Fig. 6). The ML trees grouped the isolates into three main clades (Clade I - III). One *E. coli* isolate from the ceca of chicks on reused litter made up Clade I on the core and accessory gene tree, while the number of isolates in Clade II and III strains differed between core (Fig. 6a) and accessory gene tree (Fig. 6b). Clade II isolates were clonal and recovered from chicks on fresh litter. Clade III was divided into two subclades represented by isolates from fresh (subclade IIIa) and reused litter (subclade IIIb) and subclade IIIb isolates were more clonal compared to subclade IIIa. *E. coli* Isolates classified as phylogroup A, D and F were more likely to be found in chicks on fresh litter, while phylogroup B1 was found in the ceca of chicks from reused litter (Fig. 6, Table S2). Likewise, cgMLST classification showed that ST69, ST2705 and ST6858 were associated with *E. coli* strains originating from chicks on fresh litter, while ST1403 was found in reused litter. These results suggest that the *E. coli* strains from this study have evolved to persist in either fresh or reused litter.

**Fig 6.**
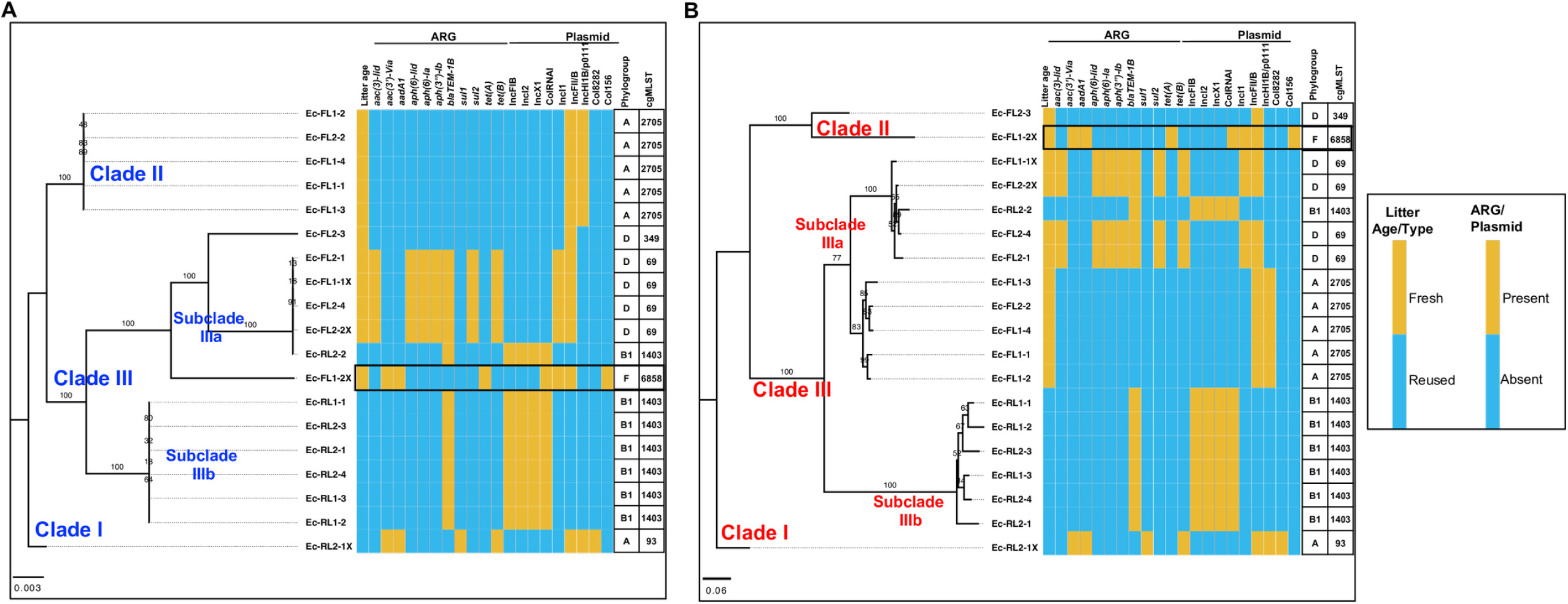
*Escherichia coli* was the main reservoir of plasmids and AMR in broiler chicks. Pan genome analysis was performed on *E. coli* strains (n =19) recovered from the ceca of broiler chicks challenged with *S*. Heidelberg and raised on fresh (FL) (n = 2) or reused litter (RL) (n=2). Maximum likelihood tree constructed using the core (**b**) and accessory **(c**) genes of the *E. coli* strains. Illumina short reads were combined with either PacBio or MinION long reads to assemble the genomes of *E. coli* strains Ec-FL1-1X, Ec-FL1-2X, Ec-FL1-3, Ec-RL2-1 and Ec-RL2-1X. GTR model of nucleotide substitution and GAMMA model of rate heterogeneity was used for sequence evolution prediction. Numbers shown next to the branches represent the percentage of replicate trees where associated isolates cluster together based on ∼100 bootstrap replicates. Clade numbers were assigned arbitrarily to ease discussion on the differences in phylogeny between isolates. Tree was rooted with Ec-RL2-1X. (Note: cgMLST – core genome Multi Locus Sequence Type). Black rectangular box shows the *E. coli* strain carrying identical IncI1-pST26 as *S*. Heidelberg.

Some *E. coli* strains from fresh litter harbored ARGs on the chromosome and plasmids. *E. coli* strains classified as pandemic lineage of extraintestinal pathogenic ST69 from fresh litter harbored a ∼128 kb genomic island encoding virulence genes (pyelonephritis-associated pili (*pap)* operon) and ARGs for aminoglycoside (*strAB* and *aph(3’)-Ia*) tetracycline (*tetB*), sulfonamide (*sul2*) and metal resistance (silver (*sil*) and copper (*pco)* operons) (Table S2; Fig. S3). The presence of a genomic island encoding AMR suggests that the ARGs are mobile. We found DNA regions encoding identical metal resistance genes (10 - 40 kb) as seen in the genomic island in p0111/IncH1B plasmids harbored by *E. coli* strains from this study and in *E. coli* genomes found in the NCBI database (Fig. S3). In addition, ST69 *E. coli* strains harbored a multireplicon IncF plasmid encoding *bla_TEM-1B_*, *aac(3)-Ild* and virulence genes (colicin M and catecholate siderophore uptake system (*iroBCDEN*)), and an untypeable IncI1 plasmid. (See Data Set S1 in the supplemental material; Data Set S1 to S5 are available in the Dryad repository online at https://doi.org/10.5061/dryad.c866t1g6c).

As previously reported [7], *E. coli* ST6858 from fresh litter carried an identical *S*. Heidelberg IncI1-pST26 plasmid, and harbored IncI2, IncF, and cryptic Col-type plasmids (Data Set S2). Antibiotic susceptible ST2705 *E. coli* isolates from fresh litter harbored no plasmid encoding antibiotic resistance genes but carried a phage-like plasmid classified as p0111 and a multireplicon IncF plasmid encoding metal resistance and virulence genes (Data Set S3). The common ARG carried by ST1403 *E. coli* strains from reused litter was *bla_TEM-_*_1B_. The gene was encoded on an IncX1 plasmid (Data Set S4), and it conferred ampicillin resistance. Additionally, ST1403 *E. coli* strains harbored IncFIB, IncI2 and ColRNAi plasmids. One *E. coli* isolate (ST93) from reused litter harbored an IncH1B/p0111 plasmid encoding ARGs for aminoglycoside (*aadA1, aac(3)- Vla*), tetracycline (*tetB*), sulfonamide (*sul1*) and mercury resistance (*mer* operon) and an IncF plasmid encoding colicin M and *iroBCDEN* genes (Data Set S5).

### Bacterial hosts from reused litter maintained IncI1 plasmid at low copies

The low rate of AMR acquisition by *S*. Heidelberg isolates from reused litter made us hypothesize that IncI1-pST26 plasmids are present at lower copies in reused litter. To test this hypothesis, we used qPCR assays targeting the region upstream of the *repA* of IncI1 plasmid including the *inc*RNAI to determine their copy number in the ceca of chicks on fresh and reused litter. We normalized the abundance of IncI1 against glyceraldehyde-3-phosphate dehydrogenase A (*gapA*) housekeeping gene encoded on the chromosome of several enteric bacteria [32] including *Salmonella* and *E. coli.* We targeted Enterobacteriaceae with *gapA* because *E. coli* was determined to be the hosts for AMR by metagenomics and WGS in this study. The relative abundance of Enterobacteriaceae bacterial population carrying IncI1 was higher in the ceca of orally challenged chicks on fresh litter compared to chicks on reused litter (W = 85, *P* = 0.00041) and was not significantly different for cloacally inoculated (W = 23, *P* = 0.24) and uninoculated control chicks (W = 63, *P* = 0.16) (Fig. 7).

**Fig 7.**
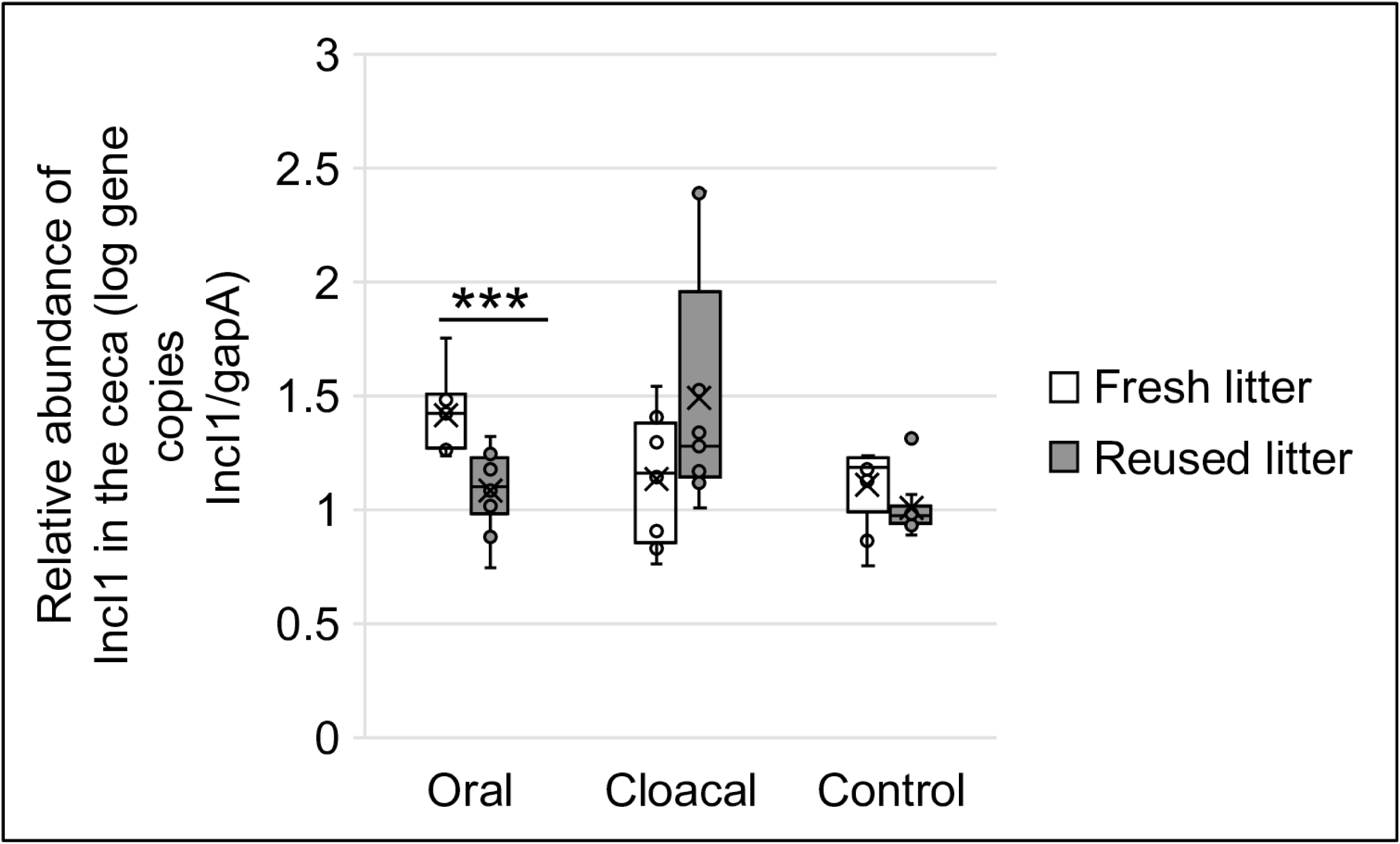
**Relative abundance of IncI1 in Enterobacteriaceae bacterial population of the ceca.** Box plot of the ratio of IncI1 gene copies/gram of ceca to gene copies of *gapA*/gram of ceca for chicks raised on fresh (oral = 9, cloacal = 8, control = 9) and reused (oral = 10, cloacal = 9, control = 10) litter. (\*\*\**P* < 0.001; Wilcoxon signed-rank test).

To further explore the effect of low copy IncI1 plasmids on AMR transfer, we selected the *S*. Heidelberg isolate from reused litter harboring IncI1-pST26 that was unexpectedly susceptible to antibiotics predicted by ARGs encoded on the plasmid. The isolate (SH-16-0-5B) was confirmed twice by broth microdilution to be susceptible to gentamicin, streptomycin, and tetracycline and once on Mueller Hinton agar supplemented with varying concentrations (0, 2, 4, 8 and 16 mg/l) of gentamicin (see supplementary methods). To ensure that the plasmid contig identified as IncI1-pST26 in this strain was not an artifact of short read sequencing, we performed qPCR on plasmid DNA and targeted four coding DNA sequence on IncI1-pST26 including its *inc*RNAi, class-1 integron (*intI1*), aminoglycoside (*aadA1*) and tetracycline resistance (*tetA*) genes (Table S3). This confirmed that the strain carried IncI1-pST26 (qPCR cycle threshold ranged from 20.2 – 27.9 for the four targets). Afterwards, we used WGS (depth of coverage of IncI1-pST26 contig/depth of coverage of the largest chromosome contig) to compare the copy number of IncI1-pST26 in SH-16-0-5B to the copy number of IncI1-pST26 in four *S*. Heidelberg isolates (hereafter referred to as SH-IncI1-FL) recovered from the ceca of cloacally inoculated chicks on fresh litter. This revealed that SH-16-0-5B maintained IncI1-pST26 at ∼0.2 copies/cell (i.e., 1 of 5 cells harbored IncI1) compared to 2.8±0.2 copies/cell in SH-IncI1-FL from fresh litter.

To investigate if the copy number difference determined using WGS exist in relevant environmental conditions, we first acclimatized the isolates in pre-reduced cecal extract (pH 6.5) for 2 h before exposing them to pre-reduced cecal extract at pH 2.5 under microaerophilic conditions. We reasoned that the low pH of the upper gastrointestinal tract (GIT) of the chicken host will interfere with the copy number of plasmids kept in a bacterial cell, which would have a direct effect on the rate of HGT. The copy number of IncI1-pST26 was significantly lower (0.00005 - 0.21 copies /per cell i.e., 1 in 20,000 to 1 in 5 cells carried IncI1) in SH-16-0-5B compared to SH-IncI1-FL (4.94 - 6.65 copies /per cell) for pH 6.5 (W = 9, *P* = 0.1) and pH 2.5 (W = 81, *P* = 4.114e-05) (Table 2). These results show that *E. coli* and *S*. Heidelberg populations in reused litter harbored lower copies of IncI1-pST26 than the populations in fresh litter.

**Table 2.** IncI1 copy number in S. Heidelberg isolates recovered from fresh and reused litter.

| S. Heidelberg Strain ID | Time (h) | pH of cecal extract | Incl1 copy number/cell<sup>a</sup>(S.D) | Coefficient of variation (%) |
| --- | --- | --- | --- | --- |
| SH-16-0-5B-RL | 2 | 6.52 | 0.001±0.001 | 155.8 |
| SH-Incl1-FL | 2 | 6.52 | 5.594±0.582 | 10.4 |
| SH-16-0-5B-RL | 0.5 | 2.52 | 0.001±0.0003 | 47.9 |
|  | 4 | 2.52 | 0.141±0.064 | 46.0 |
|  | 24 | 2.52 | 0.084±0.038 | 46.4 |
| SH-Incl1-FL | 0.5 | 2.52 | 6.304±0.165 | 2.6 |
|  | 4 | 2.52 | 6.301±0.343 | 5.4 |
|  | 24 | 2.52 | 5.892±0.378 | 6.4 |

<sup>a</sup>Three individual *S. Heidelberg* isolates from fresh litter (FL) were used to determine the copy number of Incl1 for SH-Incl1-FL, while Incl1 copy number in *S. Heidelberg* strain SH-16-0-5B from reused litter (RL) was determined using three replicates.

## Discussion

Litter is commonly used as a bedding material for raising poultry and it is ingested by chickens during pecking activities. Therefore, the microbiome of the litter is one of the first inoculum that colonizes the GIT of broiler chicks. Litter can be managed as single use i.e., complete removal of litter from a broiler house after each flock or it can be reused over multiple flocks. The benefits of litter reuse include lower cost for growing chickens and the sustainable management of litter waste [33]. It has been shown that chickens raised on reused litter harbor a different microbiome than chicks grown on fresh litter [17, 34] and that reused litter stimulates higher humoral and cell-mediated immune responses than fresh litter in chickens [35]. Muniz et al. [23] and Roll et al. [26] showed that the number of litter samples positive for *Salmonella* significantly decreases as the number of litter reuses increased compared with the first use of the litter.

In this study, we corroborate these findings by showing that neonatal chicks challenged with *S*. Heidelberg and raised on reused litter were more resistant to S. Heidelberg colonization i.e., lower positivity rate compared to chicks on fresh litter. In addition, we found that chicks raised on reused litter were less likely to be colonized with *S*. Heidelberg isolates harboring plasmids encoding AMR compared to chicks on fresh litter. We determined that the major difference between chicks grown on fresh and reused litter was the bacterial community harbored in the litter and ceca.

Pathogen induced gut microbiome modulation can create an imbalance in the abundance of bacterial species i.e., some species outgrow known resident bacteria, leading to a dysbiosis of the gut microbiome and successful *Salmonella* colonization and AMR transfer [36]. However, gut conditions that allow the maintenance of a diverse and beneficial microbiome are likely to succeed in establishing homeostasis and limit the transfer of AMR [37, 38]. In this study, we found that the microbiome of reused litter was associated with a lower HGT of AMR to *S*. Heidelberg populations in neonatal chicks. The microbiome data led us to draw one main conclusion that the litter used for growing chicks and the route of *Salmonella* challenge affected which ASVs increased or decreased in abundance in the ceca and litter.

This was the case for the ASV assigned to *Bifidobacterium* that increased in the ceca and litter of orally challenged chicks raised on reused litter but was not modulated in cloacally inoculated chicks or in chicks on fresh litter. Consequently, *Bifidobacterium* was not part of the core microbiome of the ceca or litter in this study (Table S1) and most likely evolved with neonatal chicks. *Bifidobacterium* are strict anaerobic Actinobacteria and are among the first microbes to colonize the GIT of vertebrates [39]. Additionally, *Bifidobacterium* has been shown to affect the abundance of AMR and virulence genes in the GIT of infants [40–42]. Casaburi et al [42] reported that infants fed a probiotic strain of *B. longum* harbored lower levels of ARG compared to non-fed controls. Alignment of shotgun metagenomic reads generated from the ceca of broiler chicks (n = 4) to Kraken database [43] revealed that 21% and 11% of the reads mapping to the genus *Bifidobacterium*, were aligned to *B. longum* and *B. catenulatum,* respectively (Fig. S4). Their increase in abundance for only orally challenged chicks on reused litter suggests that the route *S*. Heidelberg used for gaining entry into the GIT and the litter microbiome affected how *Bifidobacterium* was perturbed. Therefore, it is conceivable that reused litter microbiome created conditions that allowed *Bifidobacterium* to flourish and proliferate, while the microbiome in fresh litter led to a reduction or elimination of *Bifidobacterium* species.

We used metagenomics and WGS to show that *E. coli* was the main reservoir of AMR in the cecal microbiome of fresh and reused litter. Importantly, we found the identical IncI1-pST26 plasmid, acquired by *S*. Heidelberg, to be present in *E. coli*, suggesting that the plasmid was transferred in vivo within chicks raised on fresh litter. The possible sources of the *E. coli* strains in this study include the chicken host and the broiler house environment. Although, we did not establish if the meconium harbored *E. coli*, it is plausible that post-hatch chicks were colonized with *E.coli* from the hatchery [44]. The evolution of these ancestral *E. coli* strains would be shaped by selective pressures such as exposure to antibiotics and metals, and competition with resident microbiota in the environment. For chicks placed on fresh litter, the bacterial community of the first fecal droppings will compete with mostly pine shaving/plant associated microbiome. In contrast, chicks placed on reused litter will compete with microbes and metabolites deposited from earlier flocks including resident *E. coli* and *Salmonella* strains that are well-adapted to the broiler house environment. Also, the physio- chemical properties of the litter will affect the litter structure, the native microbiome and the survival of the invading pathogen. The difference in litter physio-chemical parameters in this study was inferred using pH and moisture. The pH of reused litter was higher (7.12 ± 0.26) than fresh litter (6.54 ± 0.17) (*P* = 0.028), while fresh litter had higher moisture (20.83 ± 3.54 %) than reused litter (14.65± 3.51 %) (*P* = 0.11). Litter moisture and pH are known factors that affect pathogen survival and microbial diversity [45, 46].

We did not administer antibiotics to chicks in this study and have shown that metals did not significantly affect the metabolism of *S*. Heidelberg [7]. Hence, the litter microbiome and physio-chemical properties are the major selection pressure that could explain the evolutionary trajectory of *E. coli* in this study. This hypothesis was further supported after a functional protein pathway analysis was performed on the cecal metagenome of chicks raised on fresh and reused litter. Pathways related to the biosynthesis of organic and antimicrobial molecules were enriched to a greater degree in the ceca of chicks on reused litter than chicks on fresh litter (Fig. S5). Of note was the enrichment of biosynthetic pathways for antimicrobials and secondary metabolites in reused litter (e.g., macrolides, tetracyclines, carbapenems, vancomycin and polyketides), suggesting that reused litter harbored a higher abundance of microbial species with antimicrobial properties than fresh litter.

Contrastingly, pathways associated with the biosynthesis of secondary metabolites of plants including flavonoids, indole-alkaloids and stilbenoid, diarylheptanoid and gingerol were enriched in samples from fresh litter (Fig. S5). Though we measured only two cross-sectional timepoints for reused and fresh litter, the relative abundance of plant-metabolic related pathways in fresh litter compared to the enrichment of antimicrobial molecules in reused litter suggests that a selective pressure gradient exists over time for the colonizing microbiome in reused litter. Thus, the microbiome of reused litter has the antimicrobial capability to competitively exclude invading pathogens including strains carrying AMR encoding plasmids.

Plasmid carriage is expected to impose a fitness cost on the host, but plasmid bearing microbes are pervasive in nature [47]. We showed previously that carriage of IncI1-pST26 by *S*. Heidelberg isolates from fresh litter presented a variable fitness cost to the host [7]. We found here that Enterobacteriaceae populations in the ceca of chicks on reused litter harbored lower copies of the IncI1 plasmid per cell compared to chicks on fresh litter. In addition, we did not isolate any *E. coli* isolate carrying IncI1 from the ceca of chicks raised on reused litter. The limited number of Enterobacteriaceae IncI1 donor population would affect the rate of transfer to *S*. Heidelberg recipients as direct contact is a requirement for conjugation [48]. Nevertheless, we found one *S*. Heidelberg isolate from the ceca of one chick on reused litter that harbored IncI1-pST26, however, only a fraction of its population harbored IncI1 plasmid. This isolate did not exhibit the resistance phenotype predicted by the ARGs encoded on the IncI1-pST26 plasmid. These results suggest that IncI1 plasmid carriage in reused litter posed a fitness cost on the bacterial host, but this cost was ameliorated by maintaining IncI1 at low copies in our study.

It is important to mention that the litter used for this study is not representative of all broiler litter. Litter composition may differ between farms and there are multiple physio-chemical factors that can affect the litter microbiome, e.g., moisture and pH, antibiotic usage, and feed additives. Furthermore, the age of the litter i.e., numbers of flocks that have been grown on the litter, and the length of the downtime between flocks are examples of management practices that may influence microbiome succession. Nonetheless, our study showed that the litter microbiome significantly affects the bacterial diversity of broiler chicks and the HGT of AMR.

## Materials and Methods

Details of methods used for preparing the *S*. Heidelberg inocula, challenging neonatal chicks, bacterial and DNA analyses, whole genome, and metagenome sequencing and bioinformatics have been described before for neonatal chicks raised on fresh litter [7] . We briefly redescribe these methods and present others below.

### Determining if post-hatch chicks were *Salmonella*-free

Chick pad that conveyed chicks from the hatchery were pre-enriched in 500 ml of Buffered Peptone Water (BPW) for 18-24h at 37° C. Two different enrichments broths were used to isolate *Salmonella* from the BPW pre-enrichment broths: Tetrathionate (TT; Becton-Dickinson, Sparks, MD) broth and Rappaport-Vassiliadis (RV; Becton Dickinson) media. After overnight incubation at 42°C in both enrichment broths, 10 µl aliquots from each enrichment broth was spread on two different differential media: Brilliant Green Sulfa (BGS; Becton Dickinson) agar and xylose lysine tergitol-4 (XLT-4; Becton Dickinson) agar and incubated for 18-24 h at 37° C. Isolated colonies characteristic of *Salmonella* were stabbed and streaked onto triple sugar iron agar (TSI; Becton-Dickinson) and lysine iron agar fermentation (LIA; Becton-Dickinson) and incubated for 18-24 h at 37° C for biochemical confirmation.

### *S*. Heidelberg inoculum preparation

The *S*. Heidelberg strain was made resistant to 200 ppm of nalidixic acid (nal^R^ *S*. Heidelberg) for selective enumeration and was grown overnight in poultry litter extract, centrifuged, and resuspended in 1X phosphate buffered saline (PBS). The resuspended cells were used as inocula. The complete genome of the ancestor to nal^R^ *S*. Heidelberg is available under GenBank accession number : CP066851.

### Challenging broiler chicks with nal^R^ *S*.*S*. Heidelberg

One-day-old Cobb 500 broiler chicks were either uninoculated (n =25), gavaged (n =25) or (n =25) cloacally inoculated with ∼10^6^ colony forming units (CFU) of nal^R^ *S*. Heidelberg. We also included a seeder-bird colonization method, whereby five chicks were gavaged and mingled with twenty uninoculated chicks. Afterwards, chicks were placed in floor pens at a stocking density of 0.65 m^2^/chick on fresh pine shavings (fresh litter, n =100) or reused litter (n =100). The reused litter was previously used to raise three flocks of broiler chicken’s antibiotic-free under simulated commercial poultry production conditions and was top dressed with 0.5 cm of fresh pine shavings before the placement of each flock. Broiler chicks on fresh and reused litter were housed separately and were raised for 14 days on antibiotic-free starter diet and water. At 14 days, forty-five chicks from each fresh and reused litter (ten chicks from gavaved, cloacal and uninoculated groups and fifteen chickens from the seeder method (5 seeder and 10 contact chicks) were euthanized to determine the extent of nal^R^ *S*. Heidelberg colonization in ceca. Additionally, litter samples were collected as grab samples from each pen after chicks were euthanized. The experiments were performed in April 2018. The study was approved by the University of Georgia’s Office of Animal Care and Use under Animal Use Protocol: A2017 04-028-A2.

### Determination of nal^R^ *S*. Heidelberg concentration in the ceca and litter

Ceca were removed from the eviscera of broiler chicks and stomached for 60 seconds after the addition of 3X volume to the weight (v/w) of BPW. Litter was collected as grab samples from seven locations (4 corners of the pen and 3 locations under the waterer) in each pen after chicks were removed. The litter samples were pooled and 30g was subsampled in duplicates from each pen as previously described [29]. Serial dilutions of the cecal and litter slurry were plated onto BGS containing 200-ppm nalidixic acid and plates were incubated for 24 h at 37°C.

After incubation, colonies were counted and calculated per gram of ceca or litter dry weight. When no colonies appeared, pre-enriched cecal and litter slurry was streaked onto BGS agar supplemented with nalidixic acid and incubated overnight. These plates were then examined for the presence/absence of *Salmonella* colonies. Two to six single colonies were randomly selected from each plate and archived in 30% LB glycerol at - 80 °C. In addition, cecal slurry was saved at a 4:1 ratio in Luria Bertani (LB) broth (BD Difco, MD, USA) containing 30% glycerol at –80°C, while litter samples were stored in vacuum sealed whirl pack bags at -20°C. Litter pH and moisture was determined as described previously [29].

### Antibiotic resistance phenotype determination

Antibiotic susceptibility testing was done on *S*. Heidelberg and *E. coli* isolates by following the National Antimicrobial Resistance Monitoring System (NARMS) protocol for Gram-Negative bacteria. Results were interpreted according to clinical and laboratory standards institute guidelines and breakpoints established by NARMS. In addition, we used agar dilution to determine the minimum inhibitory concentration of gentamicin for one *S*. Heidelberg isolate (SH-16-0.5B) (see supplementary methods).

### Genomic, environmental and plasmid DNA extraction

Unless otherwise noted, DNA was extracted and purified from bacterial cultures using FastDNA™ spin Kit for Soil (MP Biomedicals, LLC, CA, USA), while 250 mg of cecal slurry (previously saved in LB broth containing glycerol) and litter were extracted with Qiagen DNeasy Power-Soil DNA kit (Hilden, Germany). Extracted DNA were used for qPCR, 16S rRNA gene sequencing, shotgun metagenomics and whole genome sequencing. Plasmid DNA was extracted from two *S*. Heidelberg isolates that harbored IncI1. Isolates were cultured overnight on sheep blood agar and plasmid DNA was extracted using Qiagen Plasmid Midi kit (Qiagen Inc, Germantown, MD) as per manufacturer’s instructions. Plasmid DNA was used to construct a calibration curve for IncI1 plasmid copy number determination.

### Real-time quantitative PCR

Real-time qPCR amplification was performed as described previously (45) using a CFX96 Touch Real-Time PCR Detection System (Bio-Rad Inc., Hercules, CA). Reaction mixtures contained 1X SsoAdvanced Universal SYBR Green Supermix (Bio-Rad Inc., Hercules, CA), 600 nM (each) primers) and 2 μl of DNA. Primers used in this study are shown in Table S3. Unless otherwise stated, primers were designed with beacon designer (Premier Biosoft, Palo Alto, CA) and synthesized by Integrated DNA Technologies (Coralville, IA). Calibration curves used for converting qPCR cycle threshold values to gene copies per gram of ceca was determined using the genomic DNA from an *E. coli* strain that harbored IncI1-pST26 plasmid. To convert qPCR cycle threshold values to IncI1 copies per cell of *S*. Heidelberg, plasmid DNA of a relevant *S*. Heidelberg strain harboring IncI1-pST26 was used for calibration curve construction (Table S3). Two primer sets specific to the *gapA* gene and the region upstream of the *repA* of IncI1 including the *inc*RNAI were used to determine the gene copies of IncI1 in the ceca. The *gapA* is a housekeeping gene encoded on the chromosome of enteric bacteria including *E. coli*, *Klebsiella*, *Citrobacter* and *Salmonella* [32]. The gene copies of IncI1 per cell was determined as the copy ratio of *inc*RNAi to *gapA* [49]

### Whole genome sequencing and analysis

Whole genome sequencing libraries were prepared using either Nextera™ XT or Nextera™ DNA Flex library preparation kits (Illumina, Inc., CA, USA) following the manufacturers protocol. Libraries were sequenced on the Illumina MiSeq platform with 150 or 250-bp paired end reads. Additionally, five *E. coli* isolates were selected for long read sequencing using Sequel II System (PacBio Biosciences Inc.) or MinION device (Oxford Nanopore Technology) (File S1). Preparation and sequencing of long read libraries were done by sequencing core centers of University of Georgia and Colorado State University. The method used for read quality control and demultiplexing has been reported [7].

Genome assembly, resistome characterization, and quality assessment of assemblies was done using Reads2Resistome pipeline v.1.1.1 [50]. For tools used for genome annotation, bacteria and plasmid typing, see supplementary methods. MAFFT v. 1.4.0 [51] implemented in Geneious Prime® v 2020.0.1, was used to align and compare sequences. Roary [52] was used for pan genome analysis and phylogenetic trees were constructed using the maximum likelihood method implemented in RAxML-NG v 1.0.0 [53]. IslandViewer [54] was used to predict genomic islands.

### Identification of single nucleotide polymorphism (SNP) and Indels present in *S*. Heidelberg isolates

Alignment of raw FASTQ reads to *S*. Heidelberg (Genbank accession number: CP016573) was done using BWA (v0.7.17) [55] and SNPs/indels were called using Genome Analysis Toolkit [56]. Variant call format (VCF) files of SNPs/Indels and the script used are available in Dryad Digital Repository: https://doi.org/10.5061/dryad.c866t1g6c.

### 16S rRNA gene community analysis

The V4 hypervariable region of the 16S rRNA gene was sequenced using the paired-end (250 × 2) method on the Illumina MiSeq platform. The cecal (n = 59; 30 and 29 chicks from fresh litter and reused litter, respectively) and litter (n = 22, 11 samples each from fresh and reused litter) samples were part of a larger sequencing run and processed together with other samples. Thus, detailed sequence processing parameters are described in [7]. To avoid bias introduced by spurious amplicon sequence variants (ASVs) or from samples with low sequencing depth, any ASVs with less than 5 reads and samples with less than 5000 reads were removed from the dataset before further analysis.

Statistical analysis of microbial communities was performed in the R environment using the packages “phyloseq”, “Ampvis2”, and “vegan”. Alpha diversity indices were calculated with a dataset rarefied to the minimum sample size (8740 sequences) and compared using the Wilcoxon Signed Rank test. Principal coordinates analysis (PCoA) based on Bray-Curtis distances was performed with no initial data transformation and after removing ASVs present with relative abundance less than 0.1% in any sample. The core microbiome was determined as ASVs 0.1% relative abundance and present in at least 80 % of samples.

### Cecal metagenome sequencing

Two hundred and fifty milligrams of cecal slurry from chicks cloacally challenged with *S*. Heidelberg and raised on fresh (n = 2) or reused litter (n = 2) were used for generating shotgun and Hi-C DNA library. For shotgun library, cecal DNA was extracted with Qiagen DNeasy Power-Soil DNA kit (Hilden, Germany) and Nextera™ XT library preparation kit was used to make the library. Hi-C library was made using Phase Genomics (Seattle, WA, USA) ProxiMeta Hi-C Microbiome Kit following the manufacturer’s instructions. Libraries were sequenced by Novogene corporation (Sacramento, CA, USA) on the Illumina HiSeq platform using 150bp paired end reads. Two libraries were sequenced per HiSeq flowcell lane resulting in a total of ∼111 million shotgun reads, and ∼213 million Hi-C reads per sample. Quality controlled shotgun reads were classified using Kraken2 (v2.0.8 beta) [43] to create a count profile of the metagenome. We used cumulative sum scaling implemented in metagenomeSeq to normalize read counts [57].

### Metagenome assembled genomes and associated plasmids and ARGs

The details of the methods used for proximity-guided metagenome assembly and deconvolution have been described before [7]. Briefly, shotgun metagenomics sequence reads were filtered and trimmed for quality and normalized before assembly with metaSPAdes [58] using default options. Hi-C sequence reads were then aligned to the assembly following the ProxiMeta Hi-C kit manufacturer’s recommendations (https://phasegenomics.github.io/2019/09/19/hic-alignment-and-qc.html). Metagenome deconvolution was performed with ProxiMeta [59], resulting in the creation of putative genome and genome fragment clusters. Hi-C clusters were assessed for quality using CheckM [60] and assigned taxonomic classifications with Mash v2.1 [61] resulting in the metagenome assembled genomes (MAGs). We used PlasmidFinder (v2.1.1) [62] to identify plasmids present in MAGs.

To view AMR gene connections to hosts, AMR genes in the metagenomic assembly were annotated using NCBI’s AMRFinderPlus software (version: 3.10.5) with the --plus option specified, and all other options set to defaults. The AMR assembly annotations were then used in combination with the MAG taxonomic annotations and the mobile element to host association matrices generated by ProxiMeta to annotate which AMR genes are present in each MAG, and specify whether the gene originated from a genomic, viral or plasmid contig. This was done by making a *NxM* matrix *C*, where *N* = |*AMRa* | and *M* = |*MAG* |, where *MAG* = {set of individual metagenomic bins} and *AMRa* = {*x* ⊂ *AMR* : *x* ∈ Assembly}, where *AMR* = {set of all AMR genes in database}. The matrix (*C*) was then filled as:

For *i* in |*AMRa*|:

For *j* in |*MAG*|:

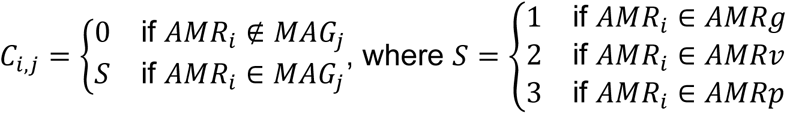

and where, *AMRg* = {*x* ⊂ *AMRa* : *x* ∈ Genomic Contigs}, *AMRv* = {*x* ⊂ *AMRa* : *x* ∈ Viral Contigs}, *AMRp* = {*x* ⊂ *AMRa* : *x* ∈ Plasmid Contigs}. The matrix, *C*, was then filtered to remove rows and columns with only 0 values and plotted using a color encoding system indicating whether the AMR gene was of genomic (cyan) or plasmid (gold) origin.

### Functional Pathway Analysis

Both shotgun sequences and ProxiMeta-deconvoluted genomes were also utilized to assess enrichment of functional pathways. Shotgun metagenomic sequences were aligned to the Hi-C genomic contigs using BWA-MEM algorithm as implemented in the BWA (v0.7.17) [55]. The Hi-C genomic contigs were then annotated using Prokka (v1.13) [63]. For each annotated region on the Hi-C genomic contigs, sequences overlapping the annotated region in the sequence alignments were counted using a custom Python script to produce a count matrix of genes for all samples. Gene counts were normalized by gene length and using total sum scaling normalization to control for differences in sequencing depth [64]. Enzyme commission identifiers were then mapped to KEGG functional pathways according to the KEGG ontology. For each pathway, an adjacency graph was reconstructed from the KEGG XML files, and connected components were determined using depth-first search [65]. Z-scores were calculated for pathways and connected components for each sample and statistical testing was then performed by comparing scores to the normal distribution. Details of this analysis is described in the supplementary methods.

### *E. coli* isolation from ceca

The method used for screening frozen cecal contents has been reported [7]. Briefly, two cecal slurry from challenged chicks raised on fresh litter and reused litter were spread plated onto CHROMagar™ plates supplemented with or without gentamicin (8 ppm) and tetracycline (8 ppm) or ampicillin (16 ppm) and cefoxitin (16 ppm) (Thermo Fisher, Waltham, MA). After 24 h incubation, blue green and blue-cream colonies were counted as presumptive *E. coli* and 4 - 5 colonies from each spread plate were picked and restruck for colony purification. No colonies grew on CHROMagar™ plates supplemented with ampicillin and cefoxitin. Colonies were preserved in LB broth containing glycerol and frozen before they were used for including solid agar competition experiment.

### Determination of plasmid copy number of IncI1

To compare the copy number of IncI1 harbored by *S*. Heidelberg strain (SH-16-0-5B) from reused litter to the copy number of IncI1 in three SH-IncI1-FL isolates from fresh litter, we exposed each isolate to acidified filter sterilized cecal extract. Cecal extract (CE) was prepared from the cecal content of 2-weeks old broiler chickens (see supplementary methods). Cecal extracts were pre-reduced by covering tubes with gas permeable paper strip and incubating overnight under microaerophilic conditions (5% O_2_, 10% CO_2_, and 85% N2) at 42°C. Three single colonies of each strain were selected from overnight cultures grown on sheep blood agar and transferred to a microcentrifuge tube containing 900 μl of pre-reduced CE (pH = 6.52), i.e., one tube per for each SH- IncI1-FL strain and three tubes for SH-16-0-5B.

After transfer, tubes were vortexed, covered with a gas permeable paper strip and incubated at 41°C under microaerophilic conditions for 2 h. After incubation, tubes were vortexed and 100 μl of the suspension was transferred to a microcentrifuge tube containing 900 μl of CE (pH of CE was adjusted to 2.52 using 1M HCl (Spectrum Chemical Mfg. Corp., CA, USA) and 1M NaOH (Fisher Chemical, NJ, USA) Afterwards, tubes were vortexed, and incubation continued for another 24 h. To determine the copy number of IncI1, one replicate tube per strain was removed at timepoints 2 h for pH 6.5 and 0.5, 4 and 24 h for pH 2.5 and used for DNA extraction. DNA was extracted from 500 - 800 μl of CE samples using FastDNA™ SPIN Kit for Soil. Calibration curves for *gapA* and *inc*RNAI were generated using genomic and plasmid DNA extracted from ancestral *S*. Heidelberg (GenBank accession number : CP066851) and one SH-IncI1-FL isolate, respectively. Plasmid copy number of IncI1 was determined as the copy ratio of *inc*RNAi to *gapA* [49]

### Statistical analyses

Continuous variables were log-transformed before any statistical tests were performed. Moreover, continuous variables did not meet the assumption of a normal distribution; therefore, non-parametric testing for direct comparisons was performed using Wilcoxon rank sum and signed rank tests, and the Kruskal-Wallis rank sum test was used for one-way analysis of variance tests. Additionally, a generalized linear model with a binomial distributed outcome and a log link function was performed as described earlier [66] to determine if there are significant differences in *Salmonella* prevalence (presence/absence) and the litter used for raising broiler chicks. The significance of the model was established using a likelihood ratio test (R function ANOVA with argument test set to “Chisq”). Multiple comparisons of means, i.e., regression coefficients for route of inoculation and litter age/type were done using the multcomp package in R. The mcp function was used to specify linear hypotheses and the glht function was used to make Tukey contrasts. Statistical analyses were performed using R (v4.0.3).

## Ethics statement

All animal experiments were approved by the University of Georgia Office of Animal Care and Use under Animal Use Protocol: A2017 04-028-A2.

## Data availability

All raw short and long FASTQ reads for *S*. Heidelberg are publicly available under NCBI accession number: PRJNA683658, while *E. coli* FASTQ reads are available under NCBI accession number: PRJNA684578. Shotgun and Hi-C reads from fresh litter and reused litter cecal samples are publicly available under NCBI accession number: PRJNA688069 and 16S rRNA gene sequences under NCBI accession number: PRJNA669215. The complete genome assemblies for *E. coli* strains Ec-FL1-1X and Ec-FL1-2X were previously published and are available under GenBank accession numbers CP066836 and JAFCXR000000000. Complete genome of *E. coli* strain Ec-RL2-1X has been made available under GenBank accession numbers CP066839. Variant call format (VCF) files of identified SNPs/indels and the Linux/Unix shell script used have been deposited in Dryad Digital Repository: https://doi.org/10.5061/dryad.c866t1g6c.

## Acknowledgements

We are grateful to Marlo Sommers, Carolina Hall, Jeromey Jackson, and Latoya Wiggins for their logistical and technical help. This work was supported by USDA Agricultural Research Service (Project Number: 6040-32000-010-00-D), non-assistance cooperative agreement (58-6040-6-030) between USDA Agricultural Research Service and University of Georgia, Research Foundation, and research service agreement (58-6040-8-035) between USDA Agricultural Research Service and Colorado State University.

This study was supported in part by resources and technical expertise from the Georgia Advanced Computing Resource Center, a partnership between the University of Georgia’s Office of the Vice President for Research and Office of the Vice President for Information Technology. Any opinions expressed in this paper are those of the authors and do not necessarily reflect the official positions and policies of the USDA or the National Science Foundation, and any mention of products or trade names does not constitute recommendation for use. IL, MOP, and JRG are employees of Phase Genomics, Inc., and have been supported in part by NIAID grants R44AI162570 and R44AI150008. The remaining authors declare no competing commercial interests in relation to the submitted work. USDA is an equal opportunity provider and employer.

## Author’s contributions

A.O., and Z.A. designed the study. A.O., G.Z., D.E.C. and C.R. performed live-broiler chicken studies. M.J.R contributed to the design of the solid agar mating experiment. A.O., J.L., G.Z., and D.C. performed bacteriological analyses, antibiotic susceptibility testing, DNA extraction and qPCR. G.Z. and A.O. performed Illumina whole genome sequencing. A.O. performed cecal shotgun and Hi-C library preparation. T.L. made 16S rRNA gene libraries and sequencing. B.Z. performed 16S rRNA bacterial community analysis and interpretation. A.O., Z.A., M.O.P., I.L., J.R.G., C.W., J.C.T., S.M.L., and R.W. performed bioinformatic analyses and data curation. A.O., J.L. and D.C. performed solid agar mating experiment. A.O., and B.Z. performed statistical analyses. A.O., Z.A., N.A.C., B.Z., R.W., M.O.P., J.R.G., S.M.L., G.Z., and J.L. drafted the manuscript, which was reviewed and edited by all authors. A.O., Z.A., and S.A.A. supervised the study.

