## Supplementary Material and Methods, Fig. S1 - S5, Table S2 and S3. for "Litter commensal bacteria can limit the horizontal gene transfer of antimicrobial resistance to *Salmonella* in chickens"

Running Title: Commensal bacteria reduce HGT

Adelumola Oladeinde[1], Zaid Abdo[2], Benjamin Zwirzitz[3¶], Reed Woyda[2¶], Steven M. Lakin[2¶], Maximilian O. Press[4¶*], Nelson A. Cox[1], Jesse C. Thomas IV[6], Torey Looft[7], Michael J. Rothrock[1], Gregory Zock[8], Jodie Plumblee Lawrence[1], Denice Cudnik[1], Casey Ritz[8] and Samuel E. Aggrey[8], Ivan Liachko[4], Jonas R. Grove[4], Crystal Wiersma[2]

**Authors' Affiliation**

[1]U.S. National Poultry Research Center, USDA-ARS, Athens, GA, USA. [2]Department of Microbiology, Immunology and Pathology, Colorado State University, Fort Collins, Colorado, USA. [3]Institute of Food Science, University of Natural Resources and Life Sciences, Vienna, Austria. [4]Phase Genomics Inc, Seattle, WA, 98109, USA. [5]Office of National Programs, USDA-ARS, Beltsville, Maryland, USA. [6]Division of STD Prevention, National Center for HIV/AIDS, Viral Hepatitis, STD and TB Prevention, Center for Disease and Control, Atlanta, Georgia, USA. [7]National Animal Disease Center, USDA-ARS, Ames, IA, USA,[8]Poultry Science Dept, University of Georgia, Athens, GA, USA

¶ These authors contributed equally to this work.

*Present address: Maximilian O. Press, Inscripta, Inc., Boulder, Colorado, USA.

### Supplementary Materials and Methods

#### Gentamicin susceptibility testing by agar dilution

Fresh Mueller Hinton II Agar (MHA) (Beckton Dickson; Franklin Lakes, NJ) supplemented with 0, 2, 4, 8 and 16 mg/L of gentamicin was prepared and used for MIC determination. To prepare bacterial inoculum for MIC determination, bacterial suspensions were made in cation adjusted MH broth (ThermoFisher; Waltham, MA) from overnight cultures of SH-16-0-5B, one SH-IncI1-FL isolate and a pan susceptible *E. coli* reference strain (ATCC 25922). The target turbidity for the suspension was a 0.5 McFarland standard. Afterwards, the viable counts in each bacterial suspension were determined by serial dilutions onto MHA and sheep blood agar (Remel; Lenexa, KS). The inoculum for the three isolates ranged from 6.2E6 cfu/ml to 1e7 cfu/ml. Three 1 μl dots and three 2 μl dots of the bacterial suspension for each isolate were placed on each of the MHA plates with gentamicin and incubated overnight at 37$^{o}$C. The lowest gentamicin concentration that inhibited growth was determined to be the MIC of the isolate.

#### Functional Pathway Analysis.

To assess enrichment of functional pathways, both the shotgun sequences and the Hi-C deconvoluted genomes were utilized for each sample. The shotgun metagenomic sequences were aligned to the Hi-C genomic contigs using the BWA-MEM algorithm as implemented in the Burrows-Wheeler Aligner software (v0.7.17) [1]. The Hi-C genomic contigs were then annotated using Prokka (v1.13) [2]. For each annotated region on the Hi-C genomic contigs, sequences overlapping the annotated region in the sequence alignments were counted using a custom Python script to produce a count matrix of genes for all samples. Gene counts were normalized by gene length and using total sum scaling normalization to control for differences in sequencing depth [3].

Enzyme Commission identifiers (EC ID) were extracted from the Prokka annotations for each gene and used to translate counts for genes into counts for each EC ID. Genes that did not have an annotated EC ID were not included in the analysis. For ambiguous EC IDs (e.g., 3.6.3.-), counts were divided evenly into the lowest-level, non-ambiguous EC IDs that were children of that EC ID. For example, if 3.6.3.- included 3.6.3.1, 3.6.3.2, and 3.6.3.3, then a count of 9 for the ambiguous EC ID 3.6.3.- would result in each of the three children EC IDs having a count of 3. EC IDs were then mapped to KEGG functional pathways according to the KEGG ontology. KEGG pathway XML files were retrieved from the KEGG database via HTTP REST query using a custom R script. For each pathway, the adjacency graph was reconstructed from the KEGG XML files, and connected components were determined using depth-first search [4]. Scores were calculated for pathways and connected components for each sample using the following equation:

$$\text{Score} = \frac{1}{K}\sum_{k=1}^{K}\frac{x_k - \bar{x}}{\sigma}$$

where $K$ is the number of nodes in each pathway or connected component, $x_k$ is the integer count for node $k$ in that pathway, $\bar{x}$ is the mean of the node counts in that pathway, and $\sigma$ is the standard deviation of the node counts in that pathway. Positive scores indicate average increased "flux" through the pathway or connected component, while negative scores indicate average decreased "flux." This approach is equivalent to averaging Z-scores for each node in each pathway or connected component. Statistical testing was then performed by comparing scores to the Z-distribution. All codes, database files, and analytic data used for functional pathway analysis are available on GitHub :

https://github.com/lakinsm/oladeinde-competitive-exclusion.

**Protein annotation and sequence typing**.

Protein annotation was done using Prokka [2], Rapid Annotation using Subsystem Technology [5] and BlastKOALA [6]. We subtyped the *E. coli* strains using ClermonTyping [7] and Multi-Locus Sequence Typing (MLST) [8], while *S*. Heidelberg was subtyped using cgMLSTFinder v. 1.1 [9]. For plasmid typing, we used plasmid MLST [10]. We confirmed that all *S*. Heidelberg isolates were *Salmonella enterica* serovar Heidelberg using *Salmonella* In Silico Typing Resource (SISTR) [11].

**Tools used for visualizing and exploring high-throughput sequence data.** BLAST Ring Image Generator [12] was used for genome comparison visualization including GC skew change and Phandango [13] was used for visualizing phylogenetic trees with their associated metadata. SnapGene® was used for drawing linear maps of plasmid, prophages, and other regions of interest in bacterial genomes.

**Cecal extraction.**

We prepared a filter-sterilized extract from the cecal contents of 2-weeks old broiler chickens raised on 3-flock old, reused litter. Ceca were aseptically removed from the eviscera of twelve broiler chicks, placed in individual stomacher bag and transported on ice to the US National Poultry Research Center for analysis. Ceca were weighed and buffered peptone water (BPW) (BD Difco, MD, USA) was added 3X volume to the weight (v/w) and stomached for 60s. Two millimeters of stomached cecal slurry was taken from each bag (n = 12) and combined. Afterwards, a 1:10 cecal slurry was made in autoclaved 1X Phosphate Buffer Saline (PBS) (Fisher Sci, Hampton, NH) and centrifuged at 4,600 x *g* for 20 min. Supernatant was sequentially filtered through 1.2 µm, 0.45 µm and 0.2 µm pore –sized polycarbonate membrane filters; and hereafter termed “cecal extract” (CE). The absence of bacteria was confirmed by culturing 100 µl of CE in Luria Bertani broth (BD Difco).

### Supplementary Table and Figure Captions

TABLE S1. List of core ASVs in ceca and litter and their taxonomy.

TABLE S2: *E. coli* genomes found in the ceca of broiler chicks.

TABLE S3. Primers used for the qPCR analysis.

FILE S1: Assembly statistics for representative *E. coli* genomes and plasmids.

**Fig S1. Maximum likelihood tree of *S*. Heidelberg isolates recovered cloacally challenged chicks**. Core genome (a) and SNP (b) based maximum likelihood (ML) tree of $nal^R$ *S*. Heidelberg isolates (n=29) recovered from the ceca and litter of chicks on fresh (n=16) and reused litter (n=13). Isolates with duplicated SNPs/indels (n=10) were removed before SNP based ML tree was constructed. All *S*. Heidelberg strains used for core genome tree were assembled using Illumina short reads except SH4-3A and SH-ancestral that were assembled using both Illumina short reads and PacBio or MinION long reads. GTR and JC model of nucleotide substitution was used for core and SNP-based tree, respectively and the GAMMA model of rate heterogeneity were used for sequence evolution prediction. Numbers shown next to the branches represent the percentage of replicate trees where associated isolates cluster together based on ~100 bootstrap replicates. SH-ancestor was used for rooting the ML trees. (Nal – nalidixic acid, Gen – gentamicin, Str – streptomycin, Tet – tetracycline, NA – Not applicable)

**Fig S2. BLASTn alignment of the IncI1-pST26 plasmid acquired by *S*. Heidelberg and IncI1-pST26** harbored by *E. coli* strain Ec-FL1-2X. IncI1-pST26 of Ec-FL1-2X was used as the reference genome.

**Fig S3. BLASTn alignment of AMR genomic island present in *E. coli* ST69 strain Ec-FL1-1X.** IncF (n = 4) and IncH1b/p0111(n =1) plasmids from this study and identical plasmids (n=3) found on NCBI were aligned against the genomic island (~128 kbp).

**Fig S4. Percentage of shotgun reads assigned to the bacterial species of the genus *Bifidobacterium.*** *Bifidobacterium* spp. read abundance were determined by aligning shotgun reads from cecal samples of chicks raised on fresh (n =2) and reused litter (n =2) to Kraken2 database. Reads were normalized by cumulative sum scaling and reads assigned to the genus *Bifidobacterium* (0.08% of the metagenomic reads) was used.

**Fig S5. Functional metabolic pathways that were enriched in the ceca of chicks from fresh litter compared to reused litter using metagenome assembled genomes.** For a pathway or connected component within a pathway (clique), the flux is the Z-score deviation from the mean taken across all samples (n = 4). Pathway scores for each connected component are shown for each sample (n = 2 for fresh and reused litter), and the difference of reused minus fresh litter scores is shown on the right column labelled "effect size. Higher effect size indicates upregulation of the pathway in the reused litter cecal samples, while lower effect size indicates upregulation in fresh litter cecal samples.

**TABLE S1 *E. coli* genomes found in the ceca of broiler chicks.**

| *E. coli* Strain ID | Litter used for raising chicks | CHROMagar (TM) plate isolate was recovered[a] | Antibiotic Resistance profile[a] | Antibiotic Resistance Genes[b] | Plasmids Incompatibility Group[c] | Phylo group | cgMLST |
|---|---|---|---|---|---|---|---|
| Ec-FL2-4 | Fresh | CHROMagar | Amp Gen Str Fis Tet | *aac(3)-Iid aph(3')-Ia aph(6)-Id aph(3'')-Ib blaTEM-1B tet (B) sul2* | IncF:B:A- IncI1 | D | 69 |
| Ec-FL2-2X | Fresh | CHROMagar + Gen + Tet | Amp Gen Str Fis Tet | *aac(3)-Iid aph(3')-Ia aph(6)-Id aph(3'')-Ib blaTEM-1B tet (B) sul2* | IncF:B:A- IncI1 | D | 69 |
| Ec-FL2-1 | Fresh | CHROMagar | Amp Gen Str Fis Tet | *aac(3)-Iid aph(3')-Ia aph(6)-Id aph(3'')-Ib blaTEM-1B tet (B) sul2* | IncF:B:A- IncI1 | D | 69 |
| Ec-FL1-1X | Fresh | CHROMagar + Gen + Tet | Amp Gen Str Fis Tet | *aac(3)-Iid aph(3')-Ia aph(6)-Id aph(3'')-Ib blaTEM-1B tet (B) sul2* | IncF:B:A- IncI1 | D | 69 |
| Ec-FL2-5X | Fresh | CHROMagar + Gen + Tet | Amp Gen Str Fis Tet | *aac(3)-Iid aph(3')-Ia aph(6)-Id aph(3'')-Ib blaTEM-1B tet (B) sul2* | IncF:B:A- IncI1 | D | 69 |
| Ec-FL2-3 | Fresh | CHROMagar | Pan susceptible | *None* | IncF:B:A- | D | 349 |
| Ec-FL2-2 | Fresh | CHROMagar | Pan susceptible | *None* | IncF:B:A- IncHI1B/p0111 | A | 2705 |
| Ec-FL1-4 | Fresh | CHROMagar | Pan susceptible | *None* | IncF:B:A- IncHI1B/p0111 | A | 2705 |
| Ec-FL1-2 | Fresh | CHROMagar | Pan susceptible | *None* | IncF:B:A- IncHI1B/p0111 | A | 2705 |
| Ec-FL1-1 | Fresh | CHROMagar | Pan susceptible | *None* | IncF:B:A- IncHI1B/p0111 | A | 2705 |
| Ec-FL1-3 | Fresh | CHROMagar | Pan susceptible | *None* | IncF:B:A- IncHI1B/p0111 | A | 2705 |
| Ec-FL1-2X[d] | Fresh | CHROMagar + Gen + Tet | Gen Str Tet | *aadA1 aac(3)-Via tet(A)* | IncF:B:A- IncI1 IncI2 Col8282 ColRNAI | F | 6858 |
| Ec-FL1-5X[d] | Fresh | CHROMagar + Gen + Tet | Gen Str Tet | *aadA1 aac(3)-Via tet(A)* | IncF:B:A- IncI1 IncI2 Col8282 ColRNAI | F | 6858 |
| Ec-RL2-1X | Reused | CHROMagar + Gen + Tet | Gen Str Fis Tet | *aadA1 aac(3)-Via tet(B) sul1* | IncF:B:A- IncHI1B/p0111 Col156 ColRNAi | A | 93 |
| Ec-RL2-4 | Reused | CHROMagar | Amp | *blaTEM-1B* | IncFIB IncI2 IncX1 ColRNAI | B1 | 1403 |
| Ec-RL2-3 | Reused | CHROMagar | Amp | *blaTEM-1B* | IncFIB IncI2 IncX1 ColRNAI | B1 | 1403 |
| Ec-RL2-2 | Reused | CHROMagar | Amp | *blaTEM-1B* | IncFIB IncI2 IncX1 ColRNAI | B1 | 1403 |
| Ec-RL1-3 | Reused | CHROMagar | Amp | *blaTEM-1B* | IncFIB IncI2 IncX1 ColRNAI | B1 | 1403 |
| Ec-RL1-2 | Reused | CHROMagar | Amp | *blaTEM-1B* | IncFIB IncI2 IncX1 ColRNAI | B1 | 1403 |

| | | | | | | | |
|---|---|---|---|---|---|---|---|
| Ec-RL1-1 | Reused | CHROMagar | Amp | *blaTEM-1B* | IncFIB IncI2 IncX1 ColRNAI | B1 | 1403 |
| Ec-RL2-1 | Reused | CHROMagar | Amp | *blaTEM-1B* | IncFIB IncI2 IncX1 ColRNAI | B1 | 1403 |
| Ec-RL1-4 | Reused | CHROMagar | Amp | *blaTEM-1B* | IncFIB IncI2 IncX1 ColRNAI | B1 | 1403 |

[a]Amp, ampicillin; Gen, gentamicin; Str, streptomycin; Tet, tetracycline; Fis, sulfisoxazole.
[b]*aph(3'')-Ib and aph(6)-Id* is the same as *StrA and StrB.*
[c]Illumina short reads were combined with either PacBio or MinION long reads to assemble the genomes of *E. coli* strains Ec-FL1-1X, Ec-FL1-2X, Ec-FL1-3, Ec-RL2-1 and Ec-RL2-1X.
[d]*E. coli* isolates carrying identical IncI1-pST26 with *S*. Heidelberg.

TABLE S3. Primers used for the qPCR analysis.

| **Target CDS** | **Target description** | **Primer Sequences (5' - 3')** | **Reference[a]** |
|---|---|---|---|
| *aadA* | Aminoglycoside resistance | F: CAGCGCAATGACATTCTTGC<br>R: GTCGGCAGCGACAYCCTTCG | 14 |
| IncI1 | *inc*RNAi | F: CAGGAGAGATGGCATGTA<br>R: GGGTTTTCCTTTTATGGC | This study |
| *intI1* | Class 1 integron | F: CCTCCCGCACGATGATC<br>R: TCCACGCATCGTCAGGC | 15 |
| *tetA* | Tetracycline resistance | F: TGTCCACCAACTTATCAG<br>R: TGCTCAGAATTACGATCA | This study |
| *gapA*[b] | Glyceraldehyde-3-phosphate dehydrogenase A (*gapA*) in *Enterobacteriaceae* | F: CCGTTGAAGTGAAAGACGGTC<br>R: AACCACTTTCTTCGCACCAGC | 16 |

[a]References 14, 15 and 16 are in supplementary material.
[b]Primers used for targeting *gapA* gene in the ceca and *S*. Heidelberg.

# A

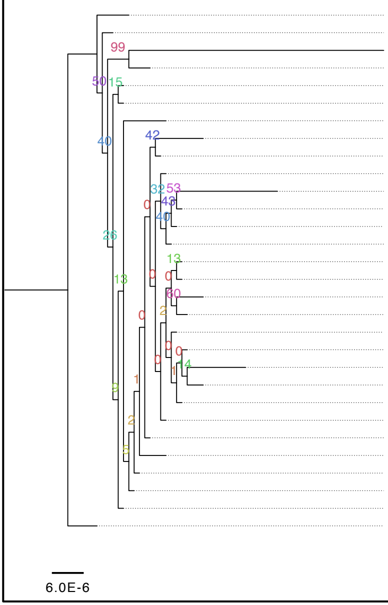


| | Isolation Source | Litter age/type | AMR phenotype | Acquired ARG | Plasmids | cgMLST |
|---|---|---|---|---|---|---|
| SH20-0-5B | Ceca | Reused | Nal | None | IncX1,Col8282, ColRNAI | 186980 |
| SH4-1A | Ceca | Fresh | Nal | None | IncX1, Col8282, ColRNAI | 186980 |
| SH20-0-5A | Ceca | Reused | Nal | None | IncX1 | 186980 |
| SH-FL1-0-5A | Litter | Fresh | Nal, Gen, Str, Tet | aac(3)-Via, aadA1, tetA | IncX1, IncI1 | 186980 |
| SH4-1B | Ceca | Fresh | Nal, Gen, Str, Tet | aac(3)-Via, aadA1, tetA | IncX1,IncI1 | 186980 |
| SH20-0-5C | Ceca | Reused | Nal | None | IncX1 | 186980 |
| SH4-3A | Ceca | Fresh | Nal, Gen, Str, Tet | aac(3)-Via, aadA1, tetA | IncX1, IncI1, Col8282 | 186980 |
| SH4-2A | Ceca | Fresh | Nal | None | IncX1 | 186980 |
| SH-FL2-0-5A | Litter | Fresh | Nal | None | IncX1 | 186980 |
| SH6-1C | Ceca | Fresh | Nal | None | IncX1 | 186980 |
| SH6-1B | Ceca | Fresh | Nal | None | IncX1 | 186980 |
| SH20-4C | Ceca | Reused | Nal | None | IncX1 | 186980 |
| SH-RL1-0-5A | Litter | Reused | Nal, Str | None | IncX1 | 186980 |
| SH16-0-5C | Ceca | Reused | Nal | None | IncX1 | 186980 |
| SH-RL2-0-5A | Litter | Reused | Nal | None | IncX1 | 186980 |
| SH-FL1-3A | Litter | Fresh | Nal | None | IncX1 | 186980 |
| SH16-0-5A | Ceca | Reused | Nal | None | IncX1 | 186980 |
| SH4-0-5B | Ceca | Fresh | Nal | None | IncX1,Col8282, ColRNAI | 171575 |
| SH16-1B | Ceca | Reused | Nal | None | IncX1 | 186980 |
| SH16-1A | Ceca | Reused | Nal | None | IncX1 | 186980 |
| SH4-0-5A | Ceca | Fresh | Nal | None | IncX1,Col8282, ColRNAI | 186980 |
| SH6-1A | Ceca | Fresh | Nal, Gen, Str, Tet | aac(3)-Via, aadA1, tetA | IncX1, IncI1, Col8282 | 186980 |
| SH6-2B | Ceca | Fresh | Nal | None | IncX1 | 186980 |
| SH6-2A | Ceca | Fresh | Nal | None | IncX1 | 186980 |
| SH20-4A | Ceca | Reused | Nal | None | IncX1 | 186980 |
| SH16-0-5B | Ceca | Reused | Nal | aac(3)-Via, aadA1, tetA | IncX1, IncI1, Col8282 | 186980 |
| SH6-2C | Ceca | Fresh | Nal | None | IncX1 | 186980 |
| SH-FL1-2A | Litter | Fresh | Nal | None | IncX1 | 186980 |
| SH-RL2-4A | Litter | Reused | Nal | None | IncX1 | 186980 |
| SH-ancestral | NA | NA | Nal | None | IncX1 | 171516 |

# B

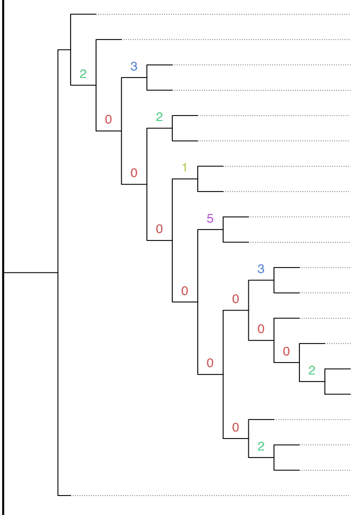

| | Litter age/type | Isolation Source | AMR phenotype | Acquired ARG | Plasmids | cgMLST |
|---|---|---|---|---|---|---|
| SH16-0-5A | Reused | Ceca | Nal | None | IncX1 | 186980 |
| SH20-0-5B | Reused | Ceca | Nal | None | IncX1,Col8282, ColRNAI | 186980 |
| SH6-1C | Fresh | Ceca | Nal | None | IncX1 | 186980 |
| SH-RL2-4A | Reused | Litter | Nal | None | IncX1 | 186980 |
| SH20-4C | Reused | Ceca | Nal | None | IncX1 | 186980 |
| SH6-2C | Fresh | Ceca | Nal | None | IncX1 | 186980 |
| SH4-1A | Fresh | Ceca | Nal | None | IncX1, Col8282, ColRNAI | 186980 |
| SH-RL1-0-5A | Reused | Litter | Nal, Str | None | IncX1 | 186980 |
| SH4-2A | Fresh | Ceca | Nal | None | IncX1 | 186980 |
| SH-FL2-0-5A | Fresh | Litter | Nal | None | IncX1 | 186980 |
| SH16-0-5C | Reused | Ceca | Nal | None | IncX1 | 186980 |
| SH6-1B | Fresh | Ceca | Nal | None | IncX1 | 186980 |
| SH4-0-5A | Fresh | Ceca | Nal | None | IncX1,Col8282, ColRNAI | 186980 |
| SH-FL1-0-5A | Fresh | Litter | Nal, Gen, Str, Tet | aac(3)-Via, aadA1, tetA | IncX1, IncI1 | 186980 |
| SH-FL1-2A | Fresh | Litter | Nal | None | IncX1 | 186980 |
| SH-RL2-0-5A | Reused | Litter | Nal | None | IncX1 | 186980 |
| SH6-2A | Fresh | Ceca | Nal | None | IncX1 | 186980 |
| SH16-0-5B | Reused | Ceca | Nal | aac(3)-Via, aadA1, tetA | IncX1, IncI1, Col8282 | 186980 |
| SH20-4A | Reused | Ceca | Nal | None | IncX1 | 186980 |
| SH-ancestral | NA | NA | Nal | None | IncX1 | 171516 |

1.0E-6

Fig. S1

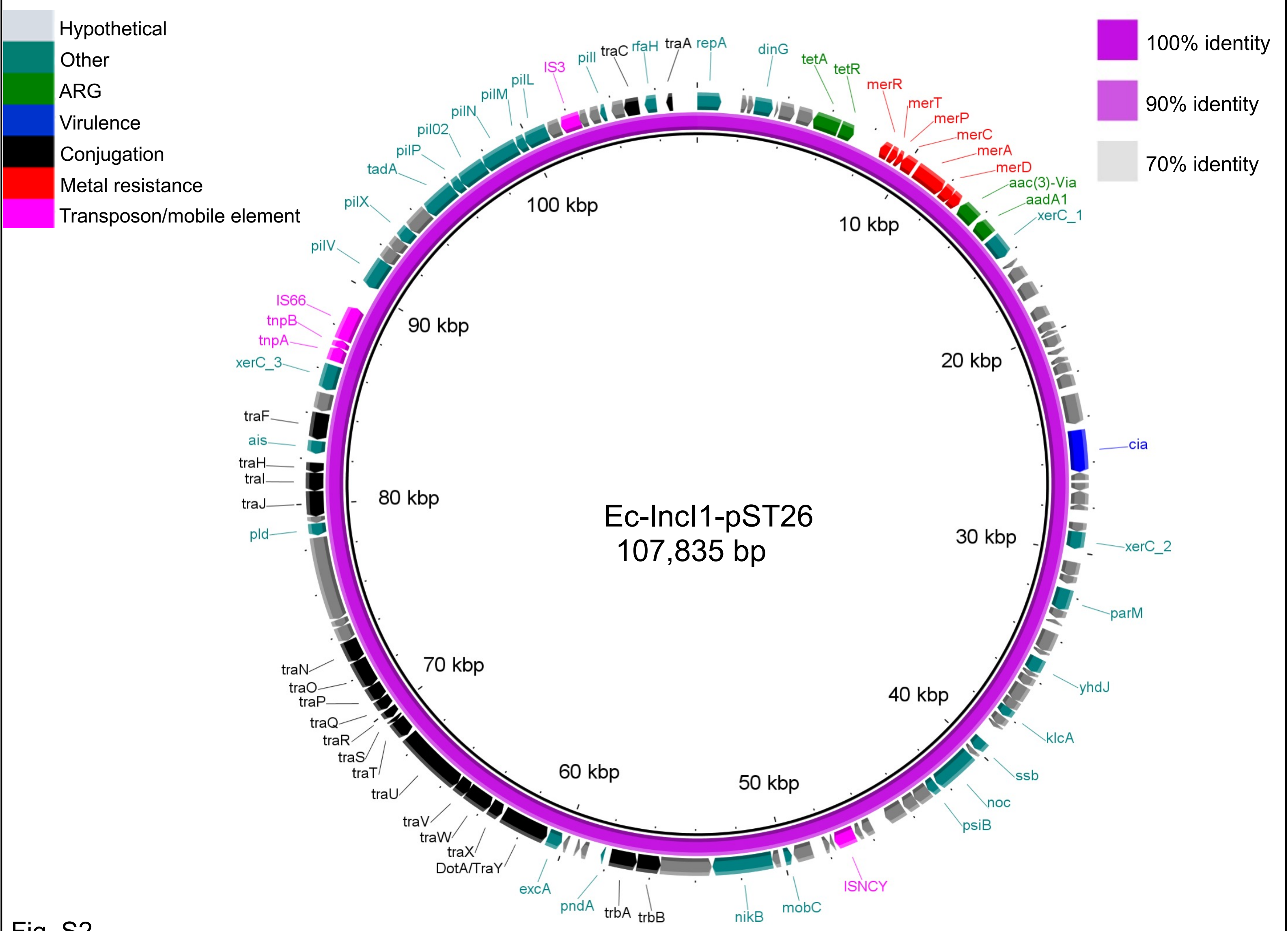
Hypothetical
Other
ARG
Virulence
Conjugation
Metal resistance
Transposon/mobile element
100% identity
90% identity
70% identity
Ec-IncI1-pST26
107,835 bp
10 kbp
20 kbp
30 kbp
40 kbp
50 kbp
60 kbp
70 kbp
80 kbp
90 kbp
100 kbp
repA
dinG
tetA
tetR
merR
merT
merP
merC
merA
merD
aac(3)-Via
aadA1
xerC_1
cia
xerC_2
parM
yhdJ
klcA
ssb
noc
psiB
ISNCY
mobC
nikB
trbB
trbA
pndA
excA
DotA/TraY
traX
traW
traV
traU
traT
traS
traR
traQ
traP
traO
traN
pld
traJ
traI
traH
ais
traF
xerC_3
tnpA
tnpB
IS66
pilV
pilX
tadA
pilP
pil02
pilN
pilM
pilL
IS3
pilI
traC
rfaH
traA

Fig. S2

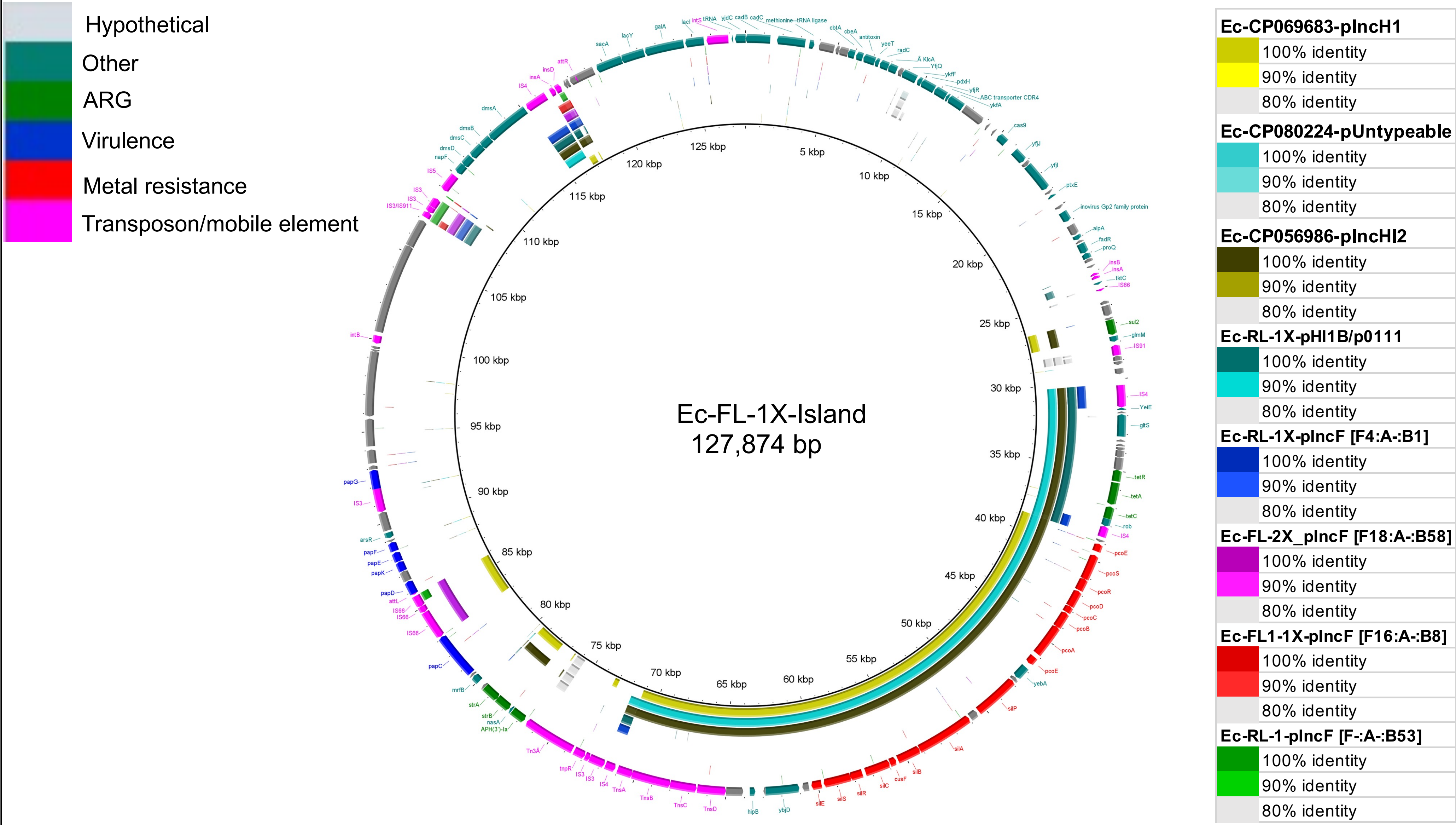
Hypothetical
Other
ARG
Virulence
Metal resistance
Transposon/mobile element
Ec-FL-1X-Island
127,874 bp
5 kbp
10 kbp
15 kbp
20 kbp
25 kbp
30 kbp
35 kbp
40 kbp
45 kbp
50 kbp
55 kbp
60 kbp
65 kbp
70 kbp
75 kbp
80 kbp
85 kbp
90 kbp
95 kbp
100 kbp
105 kbp
110 kbp
115 kbp
120 kbp
125 kbp
Ec-CP069683-pIncH1
100% identity
90% identity
80% identity
Ec-CP080224-pUntypeable
100% identity
90% identity
80% identity
Ec-CP056986-pIncHI2
100% identity
90% identity
80% identity
Ec-RL-1X-pHI1B/p0111
100% identity
90% identity
80% identity
Ec-RL-1X-pIncF [F4:A-:B1]
100% identity
90% identity
80% identity
Ec-FL-2X_pIncF [F18:A-:B58]
100% identity
90% identity
80% identity
Ec-FL1-1X-pIncF [F16:A-:B8]
100% identity
90% identity
80% identity
Ec-RL-1-pIncF [F-:A-:B53]
100% identity
90% identity
80% identity

Fig. S3

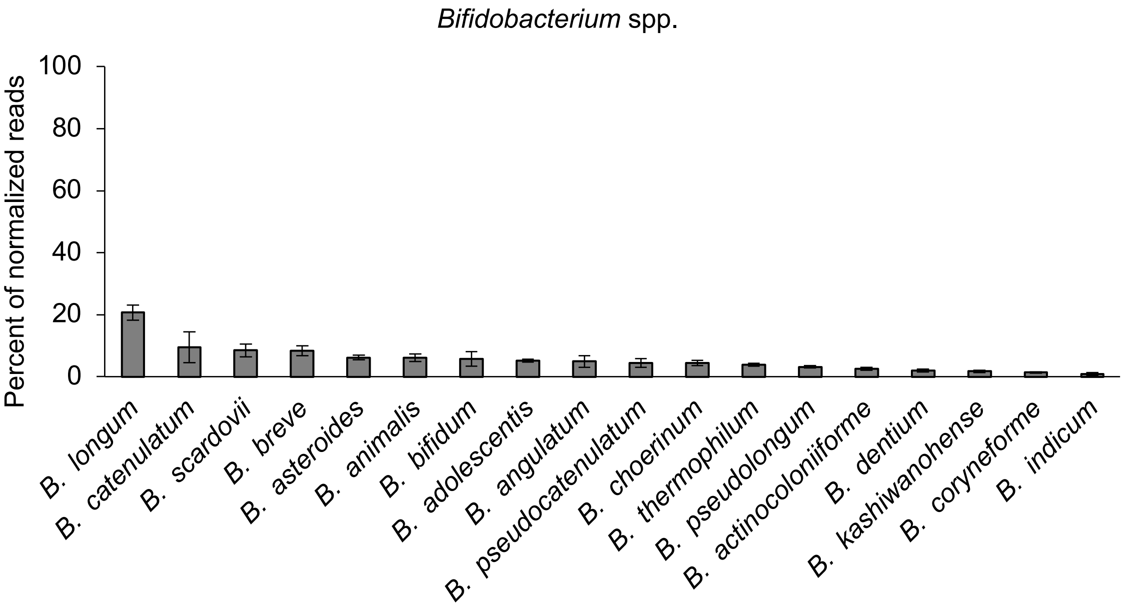

Bifidobacterium spp.
Percent of normalized reads
100
80
60
40
20
0
B. longum
B. catenulatum
B. scardovii
B. breve
B. asteroides
B. animalis
B. bifidum
B. adolescentis
B. angulatum
B. pseudocatenulatum
B. choerinum
B. thermophilum
B. pseudolongum
B. actinocoloniiforme
B. dentium
B. kashiwanohense
B. coryneforme
B. indicum


Fig. S4

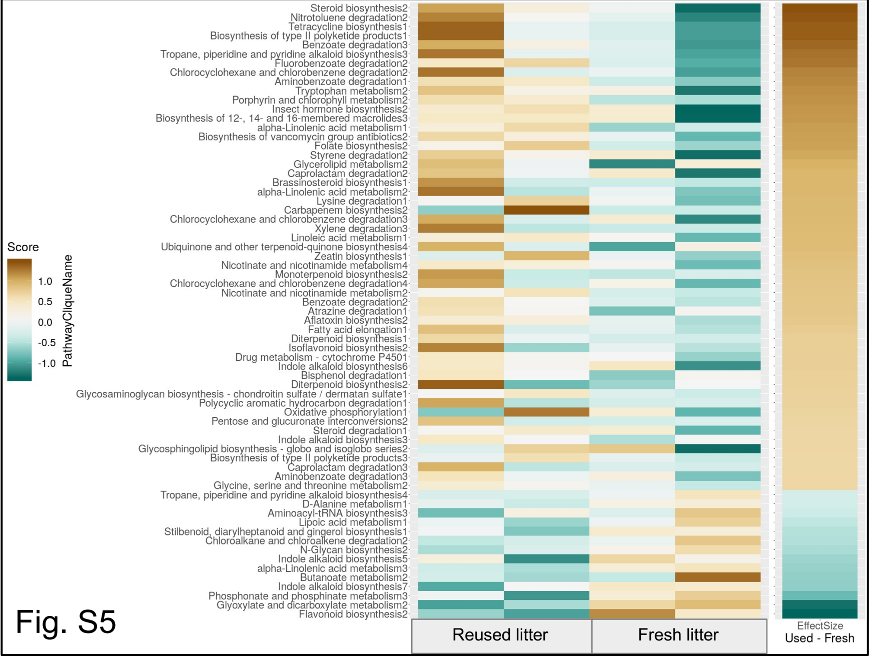
Steroid biosynthesis2
Nitrotoluene degradation2
Tetracycline biosynthesis1
Biosynthesis of type II polyketide products1
Benzoate degradation3
Tropane, piperidine and pyridine alkaloid biosynthesis3
Fluorobenzoate degradation2
Chlorocyclohexane and chlorobenzene degradation2
Aminobenzoate degradation1
Tryptophan metabolism2
Porphyrin and chlorophyll metabolism2
Insect hormone biosynthesis2
Biosynthesis of 12-, 14- and 16-membered macrolides3
alpha-Linolenic acid metabolism1
Biosynthesis of vancomycin group antibiotics2
Folate biosynthesis2
Styrene degradation2
Glycerolipid metabolism2
Caprolactam degradation2
Brassinosteroid biosynthesis1
alpha-Linolenic acid metabolism2
Lysine degradation1
Carbapenem biosynthesis2
Chlorocyclohexane and chlorobenzene degradation3
Xylene degradation3
Linoleic acid metabolism1
Ubiquinone and other terpenoid-quinone biosynthesis4
Zeatin biosynthesis1
Nicotinate and nicotinamide metabolism4
Monoterpenoid biosynthesis2
Chlorocyclohexane and chlorobenzene degradation4
Nicotinate and nicotinamide metabolism2
Benzoate degradation2
Atrazine degradation1
Aflatoxin biosynthesis2
Fatty acid elongation1
Diterpenoid biosynthesis1
Isoflavonoid biosynthesis2
Drug metabolism - cytochrome P4501
Indole alkaloid biosynthesis6
Bisphenol degradation1
Diterpenoid biosynthesis2
Glycosaminoglycan biosynthesis - chondroitin sulfate / dermatan sulfate1
Polycyclic aromatic hydrocarbon degradation1
Oxidative phosphorylation1
Pentose and glucuronate interconversions2
Steroid degradation1
Indole alkaloid biosynthesis3
Glycosphingolipid biosynthesis - globo and isoglobo series2
Biosynthesis of type II polyketide products3
Caprolactam degradation3
Aminobenzoate degradation3
Glycine, serine and threonine metabolism2
Tropane, piperidine and pyridine alkaloid biosynthesis4
D-Alanine metabolism1
Aminoacyl-tRNA biosynthesis3
Lipoic acid metabolism1
Stilbenoid, diarylheptanoid and gingerol biosynthesis1
Chloroalkane and chloroalkene degradation2
N-Glycan biosynthesis2
Indole alkaloid biosynthesis5
alpha-Linolenic acid metabolism3
Butanoate metabolism2
Indole alkaloid biosynthesis7
Phosphonate and phosphinate metabolism3
Glyoxylate and dicarboxylate metabolism2
Flavonoid biosynthesis2
Score
1.0
0.5
0.0
-0.5
-1.0
PathwayCliqueName
Reused litter
Fresh litter
EffectSize
Used - Fresh

Fig. S5
